# Double stranded RNA drives innate immune responses, sickness behavior and cognitive impairment dependent on dsRNA length, IFNAR1 expression and age

**DOI:** 10.1101/2021.01.09.426034

**Authors:** Niamh McGarry, Carol L. Murray, Sean Garvey, Abigail Wilkinson, Lucas Tortorelli, Lucy Ryan, Lorna Hayden, Daire Healy, Eadaoin. W. Griffin, Edel Hennessy, Malathy Arumugam, Donal T. Skelly, Kevin J. Mitchell, Colm Cunningham

## Abstract

Double stranded RNA is generated during viral replication. The synthetic analogue poly I:C is frequently used to mimic anti-viral innate immune responses in models of psychiatric and neurodegenerative disorders including schizophrenia, autism, Parkinson’s disease and Alzheimer’s disease. Many studies perform limited analysis of innate immunity despite these responses potentially differing as a function of dsRNA molecular weight and age. Therefore fundamental questions relevant to impacts of systemic viral infection on brain function and integrity remain. Here, we studied innate immune-inducing properties of poly I:C preparations of different lengths and responses in adult and aged mice. High molecular weight (HMW) poly I:C (1-6kb, 12 mg/kg) produced more robust sickness behavior and more robust IL-6, IFN-I and TNFα responses than poly I:C of <500 bases (low MW) preparations. This was partly overcome with higher doses of LMW (up to 80 mg/kg), but neither circulating IFNβ nor brain transcription of *Irf7* were significantly induced by LMW poly I:C, despite brain *Ifnb* transcription, suggesting that brain IFN-dependent gene expression is predominantly triggered by circulating IFNβ binding of IFNAR1. In aged animals, poly I:C induced exaggerated IL-6, IL-1β and IFN-I in the plasma and similar exaggerated brain cytokine responses. This was associated with acute working memory deficits selectively in aged mice. Thus, we demonstrate dsRNA length-, IFNAR1- and age-dependent effects on anti-viral inflammation and cognitive function. The data have implications for CNS symptoms of acute systemic viral infection such as those with SARS-CoV-2 and for models of maternal immune activation.

## Introduction

Systemic viral infection may have multiple adverse outcomes for the developing and the degenerating brain as well as producing changes in mood, behaviour and cognitive function even in normal individuals. A burgeoning field of neuroscience is now investigating several processes related to the impact of viral infection on brain function and this has become particularly pressing in the light of the SARS-CoV-2 pandemic of 2019-2021. Although some laboratories employ active infection with specific viruses, the majority studying CNS effects use the synthetic double stranded RNA (dsRNA) analogue poly I:C, since dsRNA is generated intracellularly during replication of dsRNA, positive sense RNA (+ssRNA) and DNA viruses (Weber *et al*., 2006). Poly I:C activates the immune system by signalling through toll-like receptor (Alexopoulou *et al*., 2001) and RIG-I like (RLR) receptors such as RIG-I and MDA5 (Kato *et al*., 2008), which can induce both NFκB and IRF3, with these transcription factors up-regulating production of type 1 interferons (IFN-I), other cytokines and dendritic cell maturation (Takeuchi *et al*., 2004; Stetson & Medzhitov, 2006; Gurtler & Bowie, 2013).

The behavioral and neuroinflammatory response to systemic poly I:C challenge includes hypolocomotion, reduction in species-typical behaviours and body weight and a biphasic febrile response accompanied by marked increases in peripheral and CNS pro-inflammatory cytokine expression (IFN-β, IL-6, TNF-α and IL-1β) peaking at approximately 3h post treatment (Cunningham *et al*., 2007; Konat *et al*., 2009). IL-1β is a mediator of the fever response to poly I:C in rats (Fortier *et al*., 2004) while studies using mice lacking the type I interferon receptor IFNAR1 indicate an interdependent relationship between type 1 interferons (IFN-I) and IL-6 in mediating reduced activity during sickness (Murray *et al*., 2015). More recent studies have indicated that IFNAR1, specifically in endothelial and epithelial barriers, is essential for mediating the sickness behavior response to poly I:C and that CXCL10 is a key brain mediator of these effects (Blank *et al*., 2016).

Systemically applied poly I:C has also been widely used to induce ‘maternal immune activation” in order to mimic systemic infection during pregnancy for the purposes of investigating deleterious phenotypes such as autism-like and schizophrenia-like symptoms in the offspring (Shi *et al*., 2003) and this approach has now been widely adopted (Meyer, 2014), with researchers investigating brain inflammatory, behavioural, transcriptomic and white matter changes (Garay *et al*., 2013; Crum *et al*., 2017; Richetto *et al*., 2017; Murray *et al*., 2019). Those studies aim to activate the innate immune response in a manner that mimics viral infection, but the nature of the innate immune response is often not verified, instead relying on previously published reports. However, there is significant variability in outcomes in MIA models and there is evidence from the immunology literature that responses to poly I:C are dependent on the source, length and molecular weight of the poly I:C used (Mian *et al*., 2013; Zhou *et al*., 2013). Recent studies partly characterised variable immune responses to different poly I:C preparations (Careaga *et al*., 2018; Mueller *et al*., 2019) but no study to date has examined the ability of different preparations to induce type I interferon responses despite the dominant role of these cytokines in the anti-viral response (Stetson & Medzhitov, 2006).

The importance of understanding CNS impacts of anti-viral innate immune responses has recently been given new impetus by the seismic impacts of SARS-CoV-2 on healthcare, science and society. Patients with mild symptoms include typical sickness behaviors (Eyre *et al*., 2020), while more deleterious neurological effects of the novel coronavirus have also been reported (Paterson *et al*., 2020). In particular, confusion or delirium was present in approximately 25% of all patients in a study of 20K+ patients (Docherty *et al*., 2020) and older patients more often show delirium as the initial presenting symptom (Kennedy *et al*., 2020; Poloni *et al*., 2020). The use of systemic poly I:C in older animals allows interrogation of the extent to which older individuals are more susceptible to CNS effects of systemic anti-viral responses in the absence of viable viral infection of the brain.

We hypothesized that different poly I:C preparations would produce markedly different innate immune and anti-viral inflammatory activation in periphery and brain and we thus performed a detailed analysis of innate immune responses to different dsRNA preparations. Since IFN-I is the prototypical anti-viral cytokine and an important contributor to poly I:C-induced sickness responses (Murray *et al*., 2015; Blank *et al*., 2016), we focussed particularly on the IFN-I pathway in normal and IFNAR1^-/-^ mice and also assessed the extent to which these responses were altered in aged animals and whether these lead to new cognitive sequalae. The data described herein have significant implications for understanding the CNS consequences of systemic anti-viral responses.

## Methods

### Animals

All procedures involving mice were performed in accordance with Statutory Instrument No. 566 of 2002 (Amendment of Cruelty to Animals Act, 1876) and performed, after institutional ethical review, under licence from the Department of Health and Children, Republic of Ireland and the Health Product Regulatory Authority (HPRA). C57BL/6J mice were sourced from an in-house, inbred, colony and later compared with C57BL/6J mice purchased from Harlan UK. These mice were housed in a 12h/12h light/dark cycle, in a specific pathogen free animal unit. Food and water were provided *ad libitum*.

### Treatments

Poly I:C was sourced from Sigma (Poole, UK), Amersham (GE Healthcare, Little Chalfont, UK) and Invivogen (Toulouse, France). Because the proportionate content of polymer varies between suppliers and also from lot to lot, poly I:C was initially prepared according to manufacturers instructions with equalisation of final injected concentrations but not ‘equalisation’ of stock solution concentration across different preparations. Stock solutions were as follows: Amersham (2 mg/ml), Sigma (10 mg/ml), Invivogen LMW (20 mg/ml) and Invivogen HMW (1 mg/ml), including potassium salts. Lot numbers are shown in Table 1. Poly I:C solutions, prepared with sterile saline, were mixed well and heated to 50°C (or 65-70°C in the case of Invivogen high molecular weight poly I:C) in order to solubilize and melt dsRNA and then cooled to room-temperature to allow re-annealing before freezing in 1 ml aliquots and storing at −20 °C. Concentration of double stranded poly I:C was then calculated by spectrophotometry (Nanodrop) before use and subsequently diluted to the appropriate concentration (stated as concentration of poly I:C (0.5 mg/ml), excluding salt).

**Table 1.**
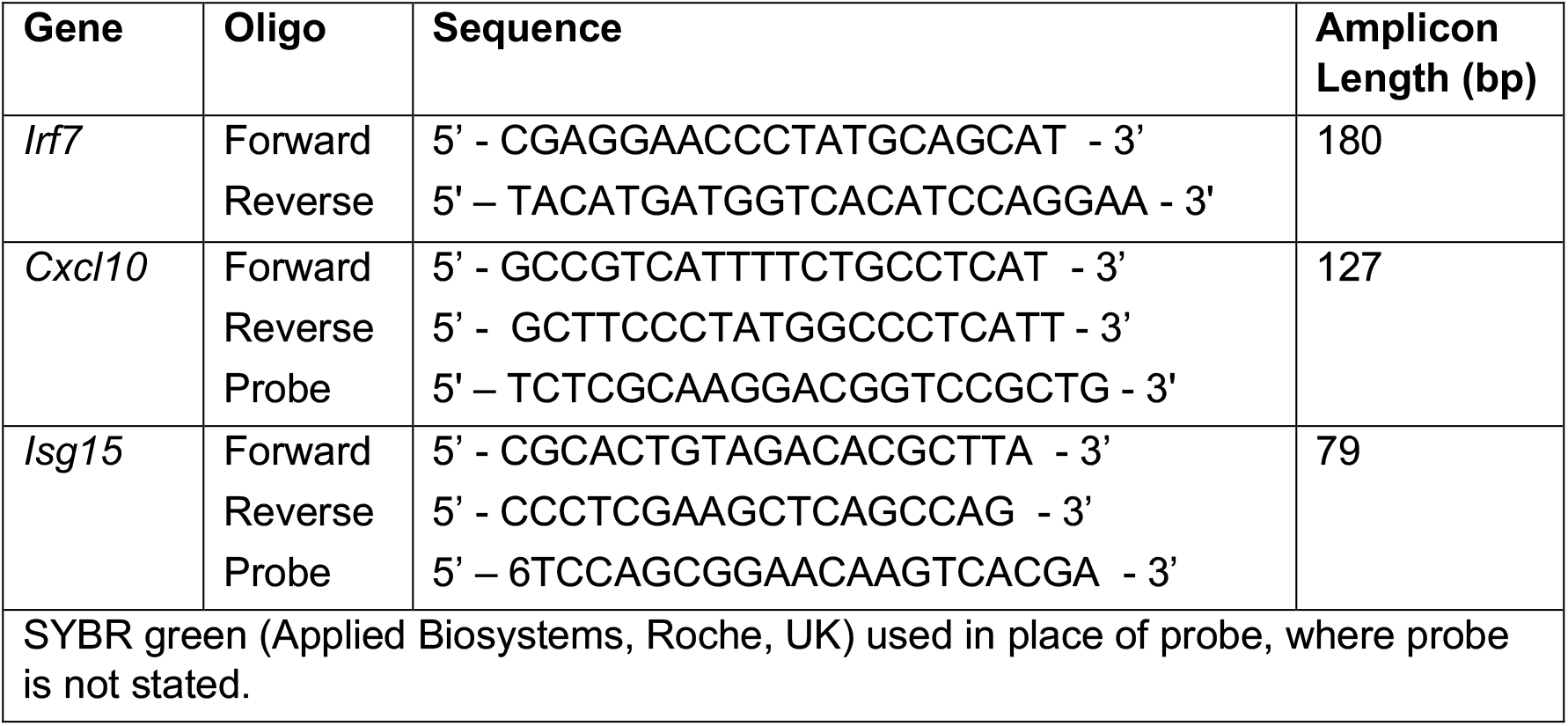

Endotoxin contamination was quantified using Limulus Amebocyte Lysate (LAL) chromogenic kit (Thermo Scientific, UK). The plate was heated to 37°C for 10 minutes (10min) following which endotoxin standards (serially diluted: 1EU/ml, 0.5EU/ml, 0.25EU/ml, 0.1EU/ml and 0EU/ml) and samples were added in duplicate (25 ul). The plates were then incubated at 37°C for 5min and the LAL lysate was added to each well (25ul). Following further 10 min incubation at 37°C the chromogenic substrate was added to each well (50ul). After 6min incubation at 37°C a 25% acetic acid stop solution was added to each well and chemi-luminescence was read at 405 nm.

### Gel Electrophoresis of poly I:C preparations

Preparations of poly I:C from different commercial sources were analysed by agarose gel electrophoresis in order to determine the RNA length i.e. the molecular weight. This included five different poly I:C preparations, two from Sigma-Aldrich (Poole, UK) and one from Amersham, of unknown molecular weight, one from Invivogen of high molecular weight (HMW) (1.5 kb – 8 kb) and one from Invivogen of low molecular weight (LMW) (0.2 kb – 1 kb). An RNA concentration was determined using A_260_/A_280_ nm using the NanoDrop 2000 spectrophotometer. Optimal parameters for gel electrophoresis encompassing a large range of RNA sizes were: 0.5% agarose gel with 15 µg of sample per lane, with visualisation using GELRED™. Agarose gel was made by mixing agarose (1 g) with 200 ml of Tris-Borate-ethylenediaminetetraacetic acid (EDTA) buffer (TBE) heating to a boil in a microwave before adding GELRED™ (15 µL), pouring and inserting the well comb. The set gel was then placed in an electrophoresis chamber, which was then filled with 1X TBE buffer and the wells loaded with amounts of between 0.1 µg and 1 µg poly I:C, with low molecular weight samples requiring loading of higher concentrations to ensure clear visualisation with GELRED™. (prepared with 4 µL loading buffer and made up to 24 µL with sterile saline). BenchTop 100bp DNA ladder (Promega) and HyperLadder™ 1000bp (Bioline) were loaded for reference. The gel was run at 100V for approximately 60 minutes, captured and analysed by exposure for 1/30^th^ using an LAS-3000 Imaging System (Fuji).

### Sickness behaviour & Working memory

#### Body temperature

Core body temperature was measured using a thermocouple rectal probe (Thermalert TH5, Physitemp, Clifton, NJ, USA). In the time-course analysis, just prior to euthanasia temperatures were recorded for each animal at 30min, 1h, 2h, 3h and 5h post-treatment with poly I:C.

#### Open field activity

Open field activity was assessed using s Benwick Electronics (AM1051) activity monitor in Perspex arena of 32 × 20 x 18 cm (height). Each activity monitor ia equipped with two sets of horizontal infra-red beams at 3 an 7 cm above the base. The beans consist of a 12 beam x 7 beam matrix, forming a grid of 66 × 2.54 cm^2^ cells and data for all beams were logged as unbroken or broken, resulting in data expressed as number of lower and number of upper beam breaks. Lower, mobile, beam breaks are used as a measure of locomotion while upper beam breaks are used to assess rears, a measure of purposeful exploratory activity. For all open field locomotor activity experiments, mice were moved from the housing room and left in the test room for 10-15 minutes prior to beginning the task to ensure an optimal state of arousal. Animals were measured for 10 minutes after placing in the middle of the open field, 3 hours post-challenge with saline or poly I:C. This time post-poly I:C, IS appropriate to capture the acute effect of poly I:C on locomotor activity (Murray *et al*., 2015). Separate cohorts of mice were tested in the open field and T maze analyses.

#### T-maze alternation: working memory

Hippocampal-dependent working memory was assessed as previously described in our “paddling” T-maze alternation task (Murray *et al*., 2015). Each mouse was placed in the start arm of the maze with one arm blocked by a guillotine style door so that they were forced to make a left (or right) turn to exit the maze, selected in a pseudorandom sequence (equal number of left and right turns, no more than 2 consecutively to the same arm). The mouse was held in a holding cage for 25 s (intra-trial interval) during which time the guillotine style door was removed, and the exit tube was switched to the alternate arm. The mouse was then replaced in the start arm and could choose either arm. Alternation from its original turn on the choice run constitutes a correct trial, meaning that 100% alternation constitutes perfect responding in this maze. On choosing incorrectly, the mice were then allowed to self-correct to find the correct exit arm. Animals were trained in blocks of 10 trials (20 min inter-trial interval) over a period of 2 weeks until reaching performance criterion: most animals reached an alternation of ≥80% but no animals were challenged with poly I:C or saline unless they showed consecutive days performing at 70% or above, showed no evidence of a side preference and did not score ≤80% in the last block of five trials before challenge. Animals were then tested 15 times over 6 hours between 3-9 hours post-poly I:C.

### Enzyme Linked Immunosorbant Assay (ELISA)

#### IL-6 & TNFα

ELISA plates (96-well; NUNC, Denmark) were incubated overnight at room temperature with capture antibody (R&D Systems, UK). Plates were washed 3 times by adding and removing 300 μl of wash buffer to each well (PBS + 0.05%Tween-20; Sigma, UK) and incubated for 1 hour with blocking buffer (Reagent Diluent, R&D Systems, UK). Plates were washed 3 times with wash buffer and 100 ul of samples and standards were added (R&D Systems, UK). Plates were left to incubate at room temperature for 2 hours. Plates were washed 3 times with wash buffer and then incubated for 1 hour at room temperature with 200 ul of detection antibody (R&D Systems, UK). Plates were washed and incubated at room temperature for 30 minutes with horseradish peroxidase-conjugated streptavidin (strep-HRP; 1:10000 Streptavidin Poly HRP, Sanquin, Amsterdam, dilution in High Performance ELISA Buffer, HPE, Sanquin, Amsterdam). Plates were washed and 100 ul of substrate solution, tetramethylbenzidine (TMB; Sigma) was added to each well. Plates were stored in the dark for 20-30 minutes or until the colour reaction developed. The colour reaction was stopped using 50 ul per well of stop solution (H_2_SO_4_ 1M). Absorbance was read at 450 nm and 570 nm using a 96-well plate reader (Labsystem Multiskan RC, UK). The readings at 570 nm were subtracted from the readings at 450 nm. A standard curve was plotted using the log of the concentrations of the standards and their relative absorbance intensities. Protein concentrations of samples were determined by reading their relative absorbance intensity from the standard curve.

#### IFNβ

ELISA plates (Biosciences, UK) came pre-coated with capture antibody. Samples and standards (100 ul) were added in duplicate, at least. Plates were left to incubate at room temperature for 1 hour. Plates were washed 3 times with wash buffer and then incubated for 1 hour at room temperature with 200ul of detection antibody (Biosciences, UK). Plates were washed and incubated at room temperature for 1 hour with horseradish peroxidase-conjugated streptavidin (strep-HRP; IFNβ: 1:285 strep-HRP dilution in Concentrate Diluent, Biosciences, UK). Plates were washed and 100 ul of substrate solution, tetramethylbenzidine (TMB; IFNβ: Biosciences, UK) was added to each well. Plates were stored in the dark for 20-30 minutes or until the colour reaction developed. The colour reaction was stopped using 50 ul per well of stop solution (H_2_SO_4_ 1 M). Absorbance was read at 450 nm and 570 nm using a 96-well plate reader (Labsystem Multiskan RC, UK). The readings at 570 nm were subtracted from the readings at 450 nm. A standard curve was plotted using the log of the concentrations of the standards and their relative absorbance intensities. Protein concentrations of samples were determined by reading their relative absorbance intensity from the standard curve.

#### IL-1β

Quantikine ELISA plates (R&D Systems, UK) came pre-coated with capture antibody. Reagent diluent (50 ul) was added to each well and samples and standards (50 ul) were added to reagent diluent in duplicate wells. Plates were left to incubate at room temperature for 2 hours. Plates were washed 5 times with wash buffer and then incubated for 1 hour at room temperature with 200 ul of IL-1β conjugate (R&D Systems, UK). Plates were washed and incubated at room temperature for 1 hour with horseradish peroxidase-conjugated streptavidin (R&D Systems, UK). Plates were washed and 100 ul of substrate solution, tetramethylbenzidine (TMB; R&D Systems, UK) was added to each well. Plates were stored in the dark for 15 minutes or until the colour reaction developed. The colour reaction was stopped using 50 ul per well of stop solution (H_2_SO_4_ 1M). Absorbance was read at 450 nm and 570 nm using a 96-well plate reader (Labsystem Multiskan RC, UK). The readings at 570 nm were subtracted from the readings at 450 nm. A standard curve was plotted using the log of the concentrations of the standards and their relative absorbance intensities. Protein concentrations of samples were determined by reading their relative absorbance intensity from the standard curve.

### Analysis of mRNA by quantitative PCR

#### RNA Extraction

Tissue was weighed and homogenised in lysis Buffer (600 μl; Qiagen RNeasy Plus Mini, RLT containing GITC RNase inhibitor) and β-mercapoethanol (10 μl/ml; Sigma, UK). Homogenised samples were filtered through Qiashredder to further homogenise tissue (Qiagen, UK) and centrifuged (11,000g; 6minutes). The Qiashredder filter units were then discarded and the supernatant was put through a genomic DNA eliminator filter and centrifuged (11,000 g; 30 seconds). The filter was discarded and 70% ethanol (350 µl; ETOH, prepared using RNase free H2O; both Sigma, UK) was added to the homogenized lysate. Qiagen RNeasy filters, which bind the RNA, were placed in new 2 ml centrifuge tubes, the lysate was loaded and the samples were centrifuged (11,000 g; 15 seconds) and the flow through was discarded. The columns were placed in new collecting tubes and RW1 Buffer was applied (350 µl; Qiagen, UK), the samples were then centrifuged (11,000 g; 15 seconds). In order to digest the DNA, rDNase Reaction Mixture (80µl; 10 µl rDNase + 70µl RDD buffer per sample; Qiagen, UK) was added directly to the centre of the silica membrane of the column and left at room temperature for 15 minutes. The silica membrane was washed by adding RW1 buffer (350 μl) and centrifuged (11,000 g; 30 seconds), and flow through discarded. RPE buffer (500 µl) was added to the filters and centrifuged (11,000 g; 15 seconds). RPE buffer (500μl) was again added to the filter and centrifuged (11,000g; 2 minutes). The column was placed in a new collecting tube and centrifuged to eliminate any carryover of RPE buffer. (11,000 g; 1 minute). The filter was placed in a new pre-labelled nuclease free 1.5 ml collection tube. The RNA was eluted using RNase free H2O (30 µl) and centrifuged (11,000 g; 1 minute) and the filters discarded. RNA concentration was quantified using the Nanodrop Spectrophotometer ND-1000.

#### Reverse transcription for cDNA Synthesis

RNA was reverse transcribed into cDNA using high-capacity cDNA archive kit (Applied Biosystems, US). The master mix was prepared from the components of the high-capacity cDNA archive kit (10X Reverse transcription buffer; 25X dNTPs; 10X RT random primers; MultiScribe reverse transcriptase (50 U/uL); RNase free H2O) the amounts of which were scaled depending on how much cDNA was needed. The Mastermix was then kept on ice. Isolated RNA was equalised to 400ng/20μl with RNase free H_2_O. Mastermix and equalised RNA were added in a 1:1 ratio. The appropriate quantity of Mastermix was added to each PCR tube and the same quantity of RNA was added. The contents were mixed by pipetting up and down and placed in the mini centrifuge to eliminate any bubbles and then placed in the thermal cycler (10 minutes at 25°C; 120 minutes 37°C; 5minutes 85°C). cDNA was stored at 4°C until needed.

#### Quantitative PCR

Mastermix was prepared using forward and reverse primers (Sigma, UK) and where possible a labelled probe for each gene (Table 1). For each gene the specific forward and reverse primers and probe were combined with Fast Start Universal Probe Master (Rox) Mastermix (Roche Diagnostics GmbH, Germany) and RNase free H_2_O (Sigma, UK). In the absence of a specific probe, Fast Start Universal SYBR Green Master (Rox) Mastermix was used (Roche Diagnostics GmbH, Germany). The finished Mastermix was added to each well of a PCR plate (24μl per well) and the cDNA was added to the Mastermix (1μl per well). The plate was run as a relative quantitative plate with the correct detectors selected using 7500 Fast System V1.3.1. The run contains 40 cycles (45 for IL-6) with the following conditions: stage 1, 50°C for 2 minutes; stage 2, 95°C for 10 minutes; stage 3, 95°C for 15 seconds and; stage 4, 60°C for 1 minute. Following the 40 cycles the plate can be removed and the data analysed using 7500 Fast system V1.3.1 relative quantitative study. A standardised control at a protein concentration of 1600ng/20ul was included on the plate in 1:4 dilutions that produced a linear amplification plot to which samples were calibrated. This control was produced using brain tissue from an animal treated IP with a high dose of the inflammatory agent LPS. This induces a strong up-regulation of inflammatory genes in all tissues. The first gene to be analysed was GAPDH and all other genes were analysed as a ratio of this genes’ expression in arbitrary units.

### Immunohistochemistry

Animals were euthanized using sodium pentobarbital (∼200μl i.p, Euthatal, Merial Animal Health, Essex, UK) and transcardially perfused with saline containing 1% heparin for 3min and then with 10% formalin for 15min to fix the tissue. The brains were then removed and post-fixed in 7ml sterilin tubes over 2 days in 10% formalin, stored thereafter in PBS and wax embedded. Brains were placed in plastic cassettes (Thermo Fisher Scientific, Dublin, Ireland) and transferred to a container of 70% alcohol for 20 minutes to begin the dehydration process. This was followed by 1.5 hourly changes in alcohol of successively increasing concentration: (70%, 80%, 95%, 100%). Cassettes were then placed in 100% alcohol III overnight. The following day cassettes went through two five hour periods in the clearing agent Histoclear II (National Diagnostics, Manville, NJ, USA) followed by a final overnight in Histoclear. The cassettes were then transferred to and put through two changes of molten paraffin wax (McCormick Scientific, St Louis, MO, USA), heated to 60°C, each for 2 hours. Embedding moulds were filled with wax, placed on a cooling stage and brains were removed from their cassettes and placed in the mould. Cassettes were placed on top of the mould and more wax was placed on top. The wax was allowed to solidify on a cooling stage. Coronal sections (10 μm) of brains were cut on a Leica RM2235 Rotary Microtome (Leica Microsystems, Wetzlar, Germany) at the level of the hippocampus and floated onto electrostatically charged slides (Menzel-Glaser, Braunschweig, Germany) and dried at 37°C overnight. Sections (10µM) of fixed tissue were dewaxed and hydrated using serial xylene, followed by histoclear and then ethanol in decreasing concentrations. The endogenous peroxidase was blocked by the quenching the tissue in 1% H_2_O_2_ Methanol and followed by antigen retrieval using citrate buffer under heat and pepsin (0.04%) digestion. Primary antibody for CXCL10 (Peprotech 1:10,000) was incubated at 4 degrees overnight in 10% normal goat serum. Secondary goat anti-rabbit (Vector, Peterborough, UK, 1:100) was incubated at room temperature as well the following signal amplifier step with ABC (Vector, 1:125). PBS wash steps were used in between each step. For final reaction the slides were placed in DAB solution and monitored by periodically checking the staining under the microscope. Finally the tissue was counterstained using haematoxylin, dehydrated and the slides were coverslipped using DPX mounting media.

### Statistical Analysis

All data are expressed as mean ± standard error of the mean (SEM) except where indicated. T-test, One-way and Two-way analysis of variance (ANOVA) tests were used to analyse data. Post-hoc analyses were performed using the Bonferroni multiple comparison test. The statistical package ‘Graph Pad Prism’ was used to complete all statistical analysis, (GraphPad Prism v9.0.0; GraphPad software, US).

## Results

### Different preparations of poly I:C have different effects on C57BL6J mice

Establishing the Maternal Immune activation (MIA) model of schizophrenia in our laboratory, using prior dose indications from other laboratories (20 mg/Kg i.p. Smith et al., 2007), we dosed pregnant C57BL6 dams with poly I:C at 20 mg/kg. Our prior characterisation of cytokine induction following systemic poly I:C treatment is one of the more widely cited accounts of innate immune activation (Cunningham *et al*., 2007) and we initially used the same poly I:C preparation that we used in those studies (i.e. that from Amersham). Using this preparation we were unable to deliver any viable offspring. Progressive reduction of the dose from 20 mg/kg to 2 mg/kg failed to prevent abortion of whole litters. Treatment, instead, with Sigma preparations of poly I:C, lead to full and healthy litters (Table 2). These preparations of poly I:C are thus likely profoundly different in their activation of innate immunity. In order to compare the potency of preparations of poly I:C from Amersham and Sigma we assessed their ability to induce sickness behaviour and pro-inflammatory cytokine responses at a dose of 12 mg/kg in adult female mice, which was that previously used in our laboratory to induce such changes. Locomotion and exploration were measured, at 2h 50min post-treatment, as the number of lower horizontal mobile beam breaks (Fig 1a) and as the number of rears, or higher horizontal beam breaks, respectively (Fig 1b).

**Table 2.** Treatment of pregnant dams with different poly I:C preparations. Avg: average; Wt: weight; Δ: change in; m: months; g: grams; M: male; F: female; * only 1 litter survived; ** weight at 3 months.

| Treatment & supplier | Lot No. | pl:C Dose, mg/kg | No. treated | $\Delta$ Wt (dam) @ 24h | Avg No.of offspring | Avg. Wt (g) offspring @ 10m: F | Avg. Wt (g) offspring @ 10m: M |
| --- | --- | --- | --- | --- | --- | --- | --- |
| Saline |  | 0 | 11 | + 3.3% | 6.6 | 25.6 | 34.6 |
| poly I:C<br>Amersham | 27-4732-01 | 20 | 1 | Died | 0 | - | - |
|  |  | 12 | 4 | -11.9% | 0 | - | - |
|  |  | 8 | 5 | -12.4% | 0 | - | - |
|  |  | 6 | 4 | -9.4% | 0 | - | - |
|  |  | 4 | 2 | -13.6% | 0 | - | - |
|  |  | 2 | 1 | -7.5% | 0 | - | - |
| poly I:C<br>Sigma | 081M4054V | 20 | 5 | +1.2% | 7 | 26.8 | 34 |
|  |  | 12 | 3 | +4.4% | 5.7 | 25.3 | 32.6 |
| poly I:C<br>Sigma | 122M4002V | 20 | 3 | +0.5% | 3.7 | 24.5* | 32.4 |
|  |  | 40 | 2 | -6.5% | 6 | 24.1** | 29.8 |

**Figure 1.**
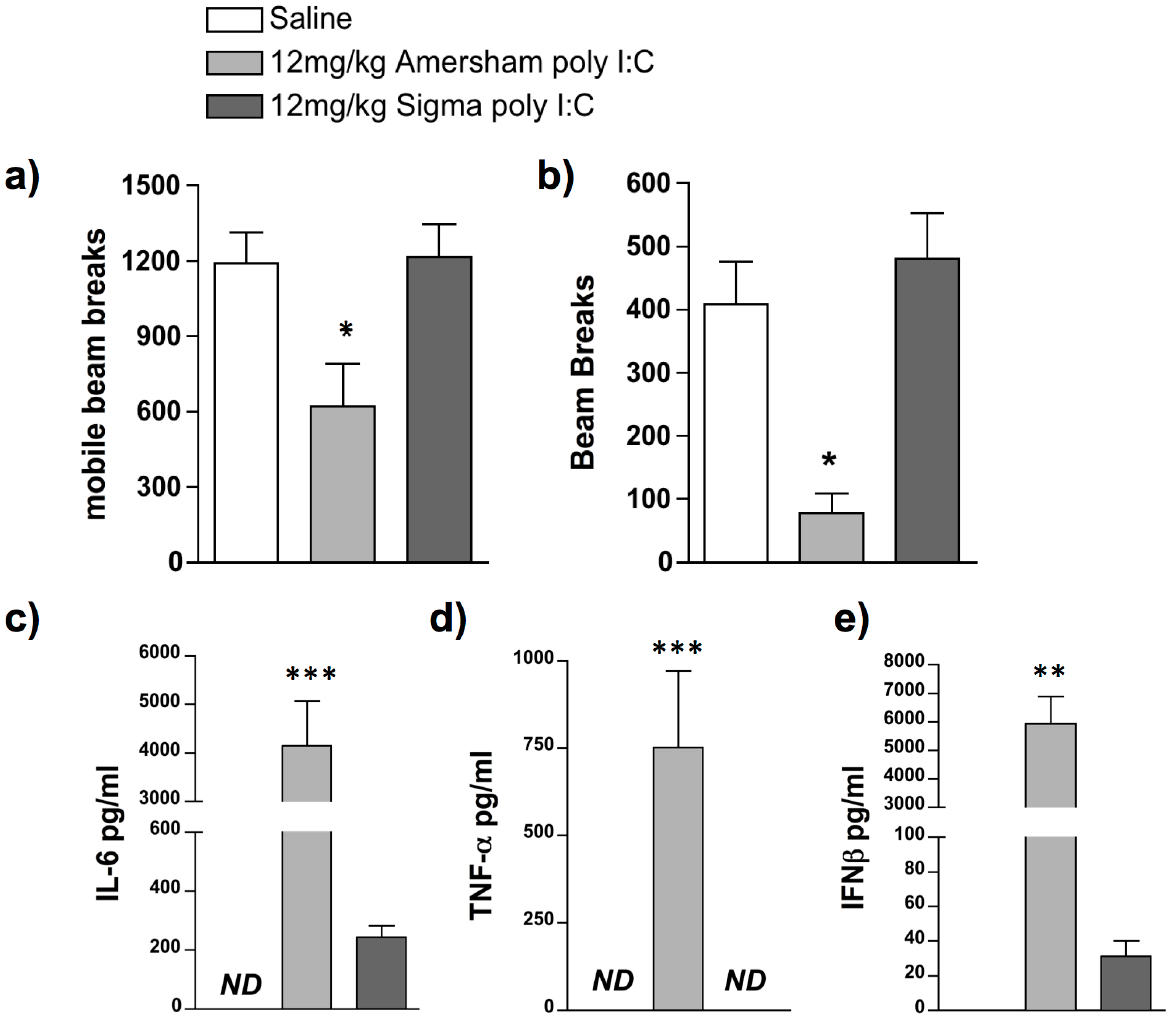
Outcomes following treatment with different poly I:C preparations. C57BL/6J female adult mice were placed in an open field 2h 50min post-treatment and measures of locomotion (a) and rears (b) were accumulated over a 10min trial period. Intraperitoneal treatments were saline (n=5), 12mg/kg Amersham poly I:C (n=4) and 12mg/kg Sigma poly I:C (n=5). **(c-e):** Systemic expression of the pro-inflammatory cytokines IL-6 (c), TNFα (d) and IFNβ (e) were determined using ELISA in C57/BL6J female adult mice 3h post-treatment in the same animals. All data are represented as mean +/− SEM and were compared using one-way ANOVA, followed by Bonferroni post-hoc tests. In a & b *p<0.05 denotes significant differences from saline-treated animals while in c-e **p<0.01 and ***p<0.001 denote significant differences between the 2 poly I:C preparations. ND denotes not-detectable.

Treatment with 12 mg/kg Amersham poly I:C caused a reduction in locomotion and exploration that is not seen in response to 12 mg/kg Sigma poly I:C. One-way ANOVA showed a significant difference between treatment groups for locomotion (F=5.368, df 2,11, p=0.0236) and exploration (F=7.899, df 2,10, p<0.0088) and Bonferroni post-hoc analysis confirmed a significant decrease in activity and rearing in animals treated with 12 mg/kg Amersham poly I:C compared to animals treated with both Saline and 12 mg/kg Sigma poly I:C, who showed no change in activity.

ELISAs were performed for IL-6 (1c), TNFα (1d), and IFNβ (1e) on plasma from blood collected at 3h post-treatment. Treatment with 12 mg/kg Amersham poly I:C produced significantly greater induction of all three pro-inflammatory cytokines than treatment with 12 mg/kg Sigma poly I:C. One-way ANOVA showed a significant difference between treatment groups for IFN-β, IL-6 and TNF-α (F≥11.67, df 2,15, p≤0.0009). Bonferroni post-hoc analysis confirmed a significant elevation in all three cytokines following treatment with 12 mg/kg Amersham Poly I:C compared to animals treated with and 12 mg/kg Sigma Poly I:C.

### Innate immune responses are determined by molecular weight of poly I:C

Hypothesising that these fundamentally different innate immune responses were underpinned by a difference in length of double stranded RNA (poly I:C) we repeated this comparison, but now included poly I:C preparations with known approximate molecular weights (Invivogen) and also assessing whether the differences in circulating cytokines were also borne out in the brain.

Levels of pro-inflammatory cytokines, IL-1β, IL-6, TNFα and IFNβ were assessed in blood plasma collected 3h post-treatment with saline, and 4 preparations of poly I:C (20mg/kg): Amersham, Sigma, Invivogen High molecular weight (HMW) and Invivogen low MW (LMW) (Fig. 2a). HMW poly I:C at doses of 20mg/kg produced systemic levels of pro-inflammatory cytokines that were much greater than those induced by low molecular weight poly I:C (Figure 2a, individual cytokines labelled therein). One-way ANOVA showed a significant difference between treatment groups for IL-1β (F=6.271, df 4,17 p=0.0027), IL-6 (F=10.96, df 4,18 p=0.0001), TNF-α (F=8.393, df 4,18 p=0.0005) and IFN-β (F=19.50, df 4,18 p<0.0001). In all cases, Amersham poly I:C was not significantly different to HMW poly I:C. IL-1β levels were significantly greater in the Amersham poly I:C and Invivogen HMW poly I:C groups compared to saline and Sigma poly I:C. Differences were even more striking for the other 3 cytokines. At 3 hours TNF-α was only detectable after Amersham and Invivogen HMW poly I:C and these treatments also induced large and comparable increases in IFN-β that were 30-100 fold higher than those present in Sigma and LMW Invivogen poly I:C-treated animals. IL-6 levels were 10-fold greater in Amersham vs. Sigma preparations and 4 fold greater in HMW vs. LMW (Bonferroni post-hoc analysis in figure legend 2).

**Figure 2.**
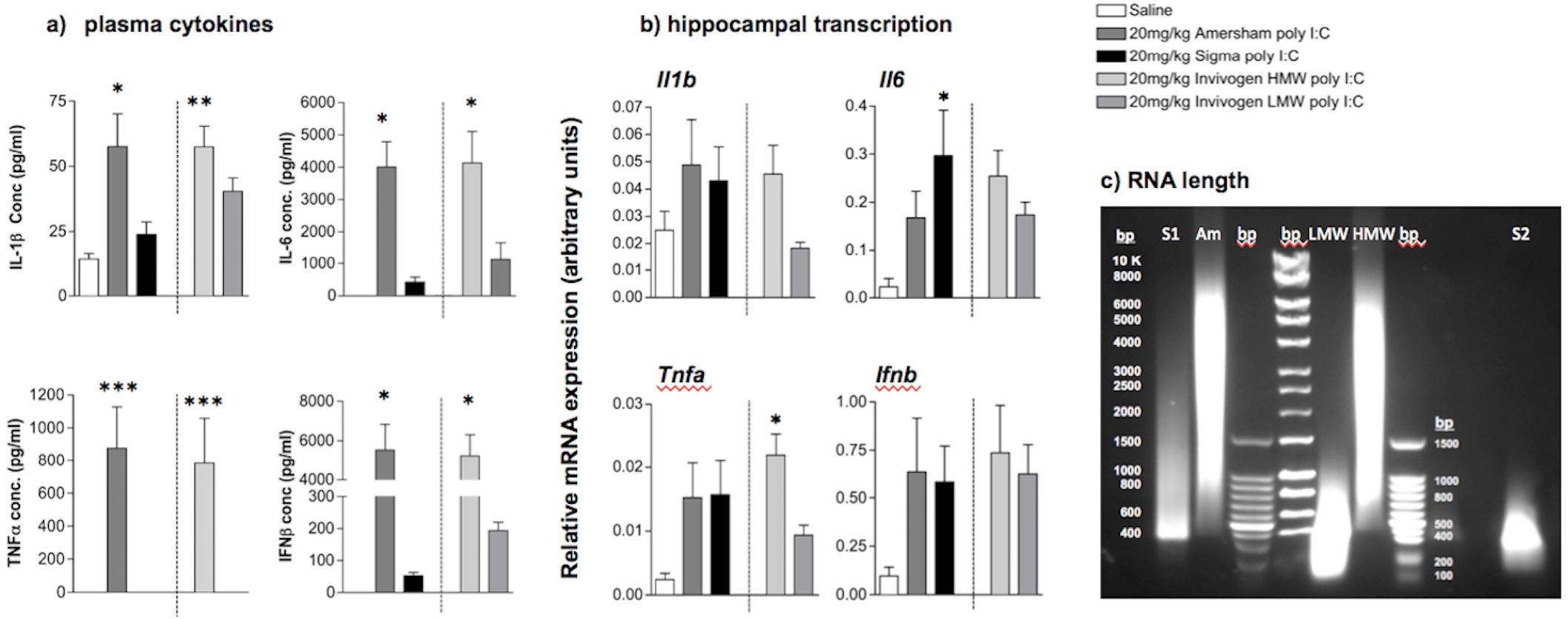
Comparison of poly I:C preparations. **(a)** Plasma concentration of the pro-inflammatory cytokines IL-1β, IL-6, TNFα, IFNβ were determined using ELISA. **(b):** Transcription of the pro-inflammatory cytokines IL-1β, IL-6, TNFα, IFNβ in hippocampal (HPC) tissue were determined using real time PCR. Experimental groups consisted of in C57BL/6J female adult mice 3h post-treatment with saline (n=3), 20mg/kg Amersham poly I:C (n=5), 20mg/kg Sigma poly I:C (n=5), 20mg/kg HMW Invivogen poly I:C (n=5) and LMW Invivogen poly I:C (n=5). (c) 0.5% agarose gel showing approximate length of different preparations in base pairs (bp). S1, S2 represent 2 different batches of Sigma polyI:C; Am represents Amersham poly I:C; LMW: In vivogen low molecular weight, HMW: Invivogen high molecular weight. All data are represented as mean +/− SEM and were compared using one-way ANOVA, followed by Bonferroni post-hoc tests. For plasma cytokines, Bonferroni tests compared differences between Sigma poly I:C versus all other poly I:C groups, and differences are denoted by *p<0.05, **p<0.01 and ***p<0.001. For hippocampal transcripts increases were smaller and more variable so Bonferroni tests were performed to compare different poly I:C treatments to saline-treated animals and differences are denoted by *p<0.05.

Transcripts for inflammatory cytokines in the hippocampus were assessed in the same animals to compare peripheral and central responses (Fig. 2b). Quantitative PCR was performed for *Il1b, Il6, Tnfa* and *Ifnb* on cDNA synthesized from RNA isolated from dissected hippocampus. All poly I:C preparations produced increases in mRNA induction for *Il6, Tnfa* and *Ifnb* but variable and limited induction of *Il1b* mRNA was seen. The large inductions seen in the plasma, that differed markedly dependent by different poly I:C preparations, were not obvious in the hippocampus. One-way ANOVA showed a significant difference between treatment groups for *Il6* (F=3.455, df 4,20, p=0.0266) and *Tnfa* (F=4.153, df 4,19, p=0.0139), no significant differences were seen for *Il1b* (F=1.674, df 4,18, p=0.1997). Thus, despite wide divergence between HMW and LMW poly I:C in the production of systemic cytokine synthesis, all cytokines were approximately equal in the hippocampus following treatment with all 4 poly I:C preparations.

The different inflammatory potencies observed with Amersham and Sigma poly I:C preparations, appeared to align to with the differences between HMW and LMW preparations. Therefore poly I:C preparations were run in a 0.5% agarose gel to assess their molecular weights. This agarose gel comparison (Fig. 2c) confirmed the predicted differences in molecular weight: Amersham poly I:C ran as a smear between 1000 and 6000 bp, centred on approximately 5000 bp, very similar to that observed for invivogen HMW. Two different preparations of Sigma poly I:C ran at approximately 400 bp and 300 bp, similar to that shown by Invivogen LMW. This confirms that Sigma and Amersham preparations of poly I:C are fundamentally different in length: Amersham is HMW and at least these two Sigma preparations are LMW. We also assessed endotoxin contamination using LAL Chromogenic Endotoxin Quantitation. Sigma poly I:C showed detectable levels of endotoxin contamination (36 EU/ml) while all other samples showed negligible levels (0.4-0.6 EU/ml; data not shown). Therefore endotoxin contamination does not contribute meaningfully to differential potency of different preparations used here.

### Robust Sickness behaviour but limited IFNβ is achieved at higher doses of LMW poly I:C

Since Sigma poly I:C was ineffective at inducing sickness behaviour at 12 mg/kg and at producing TNF-α and IFNβ at 20 mg/kg we assessed whether this relative unresponsiveness to LMW poly I:C could be overcome at significantly higher poly I:C doses. We challenged animals at 40 and 80 mg/kg poly I:C (Sigma) and compared these to 20 mg/kg poly I:C from Amersham, assessing sickness behaviour, circulating IL-6, TNFα, and IFN-β and liver transcription of the same cytokines. LMW poly I:C induces sickness behaviour responses that are even more robust than HMW poly I:C if administered at sufficiently high doses (Fig. 3a). All poly I:C treatments produced significant decreases in locomotor activity and rearing (after a significant one-way ANOVA, all treatments were p<0.001 vs saline by Bonferroni post-hoc tests). Moreover, LMW at 40 mg/kg and 80 mg/kg produced suppression of locomotion and rearing activity that was significantly greater than Amersham poly I:C at 20 mg/kg (p<0.01 for activity and p<0.05 for rears).

**Figure 3.**
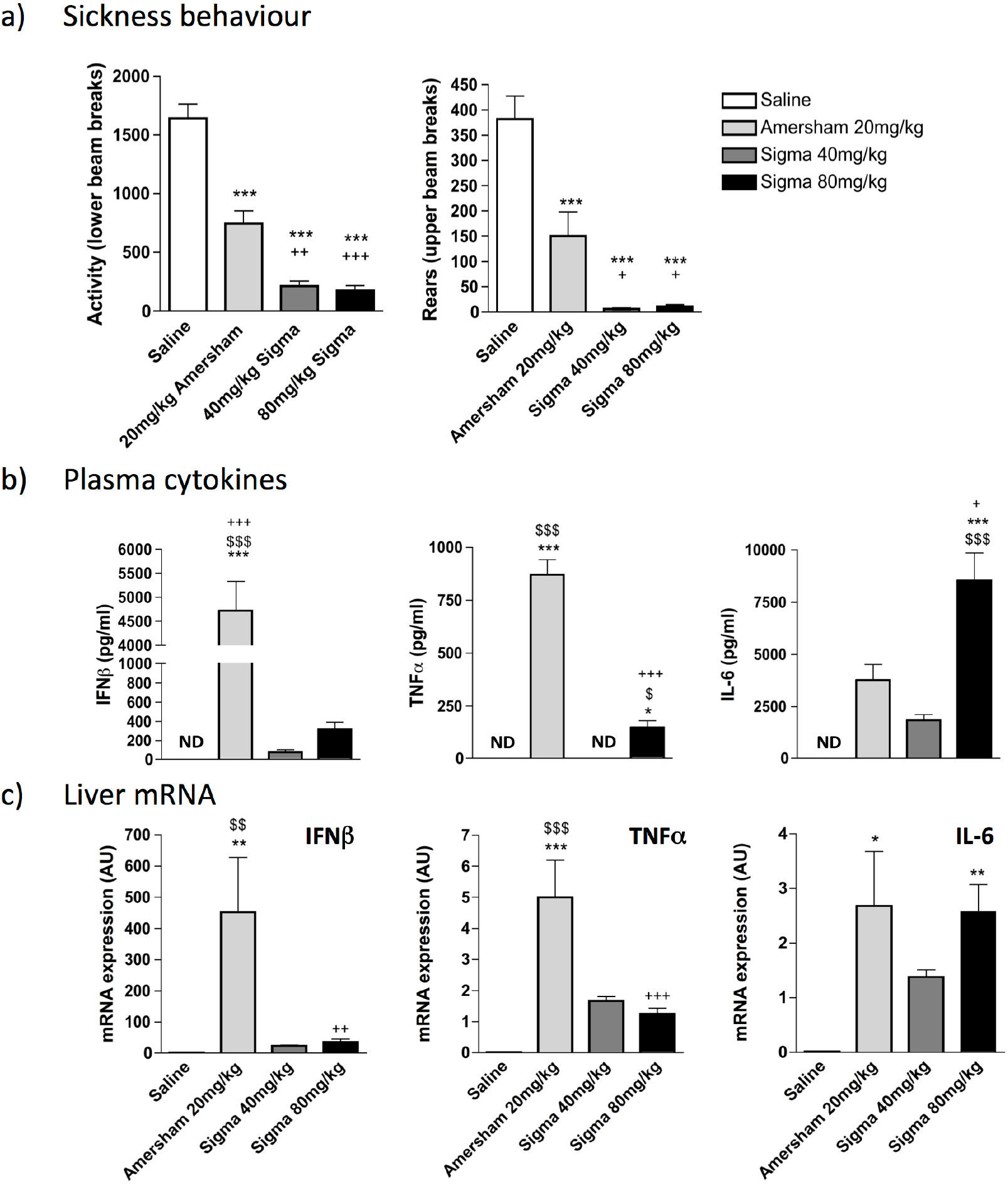
Effects of increasing the dose of LMW poly I:C. **a)** C57BL/6J female adult mice were placed in an open field 2h 50m post-treatment and measures of locomotion (lower beam breaks) and rears (upper beam breaks) were accumulated over a 10m trial period. **b)** Systemic expression of the pro-inflammatory cytokines IL-6, IFNβ and TNFα were determined using ELISA. **c)** Transcription of the pro-inflammatory cytokines *Il6, Ifnb* and *Tnfa* in liver tissue were determined using real time PCR. Data represent mean +/− SEM for saline (n=4), 20mg/kg Amersham poly I:C (n=4), 40mg/kg Sigma poly I:C (n=5) and 80mg/kg Sigma poly I:C (n=5). All data were compared using one-way ANOVA, followed by Bonferroni post-hoc tests comparing differences between each treatment group. *p<0.05, **p<0.01 and ***p<0.001 denote significant differences between saline and the represented poly I:C preparation. ^$^p<0.05, ^$$^p<0.01 and ^$$$^p<0.001 denote significant differences between 40mg/kg Sigma poly I:C and the represented poly I:C preparation. ^+^p<0.05, ^++^p<0.01 and ^+++^p<0.001 denote significant differences between 20mg/kg Amersham poly I:C and the represented poly I:C preparation.

Analysis of circulating cytokines (Fig. 3b) showed that levels of IFNβ remain at very low levels even with these highly sickness-inducing doses of Sigma poly I:C. After a one-way ANOVA (F=20.75, df 3,13 p<0.0001) only Amersham poly I:C showed significant elevation of IFNβ and levels were more than 15 fold higher than those induced by 80 mg/kg Sigma poly I:C. TNFα showed a similar trend (F=146.7, df 3,13, p<0.0001) with very high levels observed only with Amersham poly I:C, although levels induced by 80 mg/kg Sigma were now statistically significantly different from saline, albeit still 8 fold lower than Amersham poly I:C.

IL-6 was quite distinct from this profile (F=146.7, df 3,13, p<0.0001). There was a clear dose-response relationship between Sigma poly I:C dose and circulating IL-6 with levels after 80 mg/kg now statistically significantly higher than all other groups (p<0.05 with respect to Amersham poly I:C; Figure 3b).

Comparing these levels to liver transcription of the genes for these cytokines proved quite consistent with the circulating measures. Similarly to the systemic response 20mg/kg Amersham poly I:C induced robust induction of IFN-β and TNF-α mRNA that was not seen following treatment with either 40 mg/kg or 80 mg/kg Sigma poly I:C (Figure 3c). One-way ANOVA showed significant differences between treatment groups for IFN-β (F=10.60, df 3,13, p=0.0008) and TNF-α (F=19.81, df 3,13, p<0.0001) and Bonferroni post-hoc analysis showed that Amersham>Sigma 80 mg/kg (p<0.01 for IFNβ and p<0.001 for TNFα). However, the 80 mg/kg dose of Sigma poly I:C induced very robust increases in hepatic IL-6 mRNA transcription, which were not significantly different from those induced by Amersham poly I:C (F=7.032, df 3,13, p<0.0047). The pattern of pro-inflammatory cytokine induction seen in the blood is quite well reflected in liver pro-inflammatory cytokine gene transcription.

### Time course analysis of innate responses to poly I:C of different MWs

Given the robust sickness responses to high dose LMW poly I:C we sought to assess whether the 3h time point used for previous analysis of innate immune responses might have missed increases in some analytes with LMW poly I:C. Therefore we compared 80 mg/kg LMW poly I:C to 20 mg/kg HMW poly I:C at 30, 60, 120, 180 and 300 minutes post-challenge.

a) Plasma cytokines. All cytokines were increased in time-dependent fashion (main effects of time and treatment by one way ANOVA analysis for all analytes) and we present selected Bonferroni post-hoc results. IL-6 was significantly higher with LMW poly I:C but shared a very similar time course with HMW, barely increasing at 60 minutes and peaking at 120 minutes but remaining high at 180 minutes before largely resolving by 300 minutes (LMW > HMW at 120 & 180 min, Bonferroni p<0.0001 ****). TNF-α showed a different time course: LMW poly I:C TNF-α was markedly induced by 60 min, while secretion had not yet begun with HMW (p<0.01 +). Both LMW and HMW had high expression at 120 min (both +++ vs saline) but TNF secretion had resolved by 180 mins after LMW but remained highly elevated 180 min post-HMW poly I:C (HMW > saline p<0.01 ++ and > LMW *). Resolution occured by 300 min post-HMW poly I:C. With HMW poly I:C IFNβ showed no induction at 30 or 60 min, very strong induction at 120 min (P<0.0001) which increased further at 180 min (P<0.0001) before partial resolution at 300 min. Conversely, LMW induced very little IFNβ, with a small increase at 120 min (P<0.05 vs. saline +) and essentially nothing thereafter.

b) Hypothalamic transcripts. The timecourses of *Il6, Tnfa, Il1b* were very similar between HMW and LMW poly I:C, although levels of *Il6* were much higher after LMW. Comparing transcription of *Ifnb* and interferon-responsive genes revealed a remarkable pattern. Although the expression of *Ifnb* was apparent in both groups by 180 min, it was much higher by 300 min after LMW compared to HMW (p< 0.0001). However the expression of the IFN-dependent genes *Irf7* and *Eif2ak2* (PKR) was very much higher post-treatment with HMW poly I:C. This suggests that the induction of interferon-dependent genes in the hypothalamus is driven by circulating IFNβ (Figure 4a) rather than by local, hypothalamic synthesis of IFNβ (Figure 4b).

**Figure 4.**
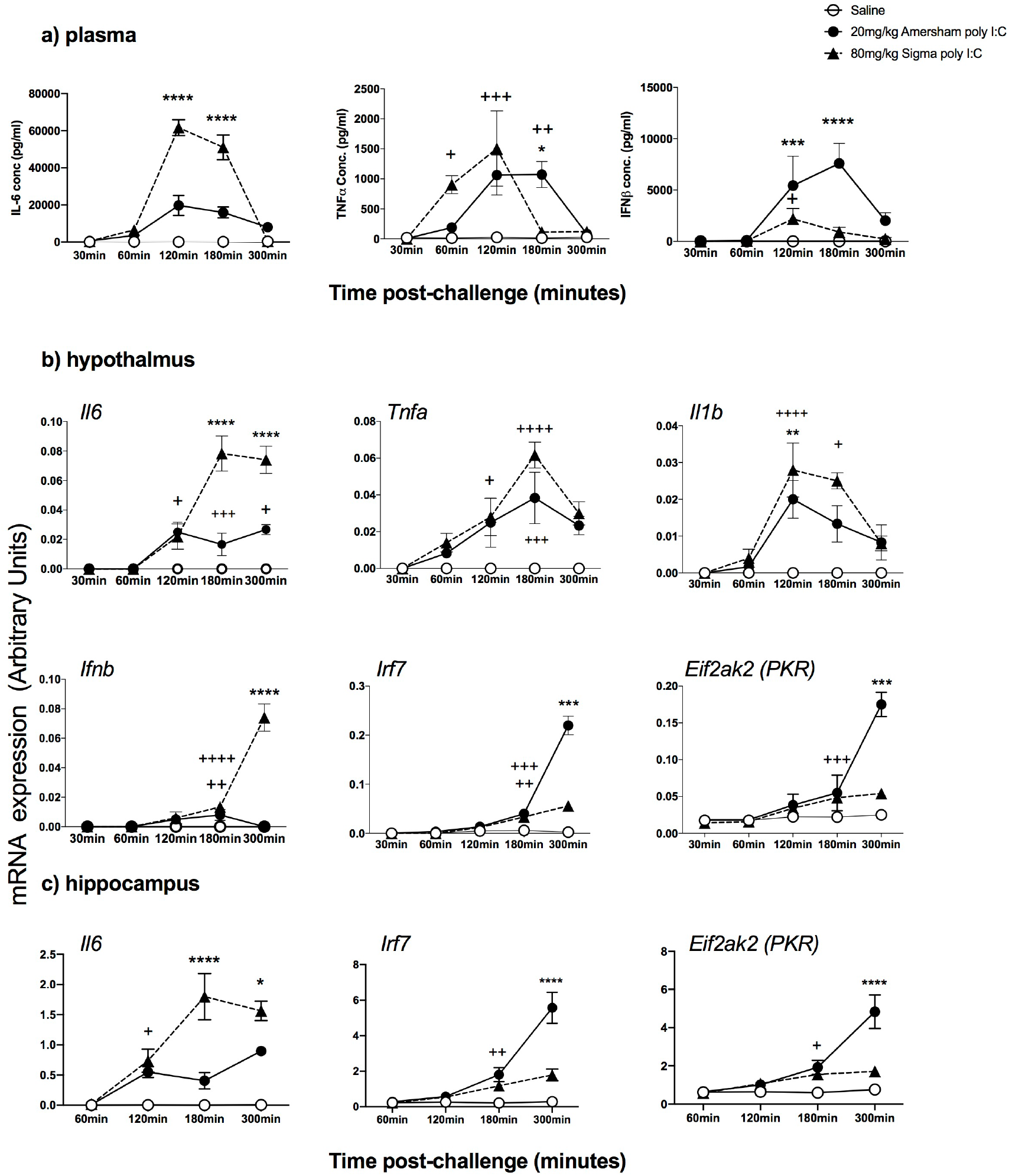
Time course analysis of innate responses to poly I:C of different MWs. a) Systemic secretion of the cytokines IL-6, TNFα and IFNβ was determined using ELISA in female adult mice following treatment with saline (n=4 per time-point), 20mg/kg (n=6 per time-point) and 80mg/kg Sigma poly I:C (n=5-6 per time-point). b) The same animals were used for isolation of hypothalamic RNA for cDNA synthesis and qPCR for expression of key pro-inflammatory transcripts and type I interferon pathway transcripts. c) The same animals were used for isolation of hippocampal RNA to verify these patterns in hippocampus. All data are represented as mean ±SEM and were compared using two-way ANOVA, followed by Bonferroni post-hoc tests comparing differences between saline and both poly I:C groups (^+^ p<0.05, ^++^ p<0.01 and ^+++^ p<0.001) and between the 2 different poly I:C preparations (*p<0.05, **p<0.01, ***p<0.001, ****p<0.0001).

A similar pattern of inducing significantly higher *Il6* witih LMW but much higher *Irf7* and *Eif2ak2* (PKR) with HMW poly I:C was also observed in the hippocampus (Figure 4c) indicating that in multiple brain regions systemic LMW poly I:C induces higher brain transcription of innate genes but downstream, interferon-dependent genes are actually driven by circulating IFNβ.

#### Relationship between circulating IFNβ and brain interferon-dependent responses

Given the finding that brain expression of IFN I-dependent genes associates strongly with circulating IFNβ rather than brain expression of *Ifnb* transcripts, we assessed an additional panel of IFN-I-dependent genes to assess whether circulating IFNβ was sufficient to induce their expression in brain. In order to examine whether their expression was exclusively regulated by IFNβ, we assessed whether IFNAR1 was necessary to facilitate their expression upon poly I:C treatment. IFNβ, administered at a dose of 25,000 units, induced robust increases of *Irf7, Isg15* and *Cxcl10* in the hippocampus of normal C57BL6J mice with respect to saline challenges in the same animals (Figure 5a, all p<0.01 by students t-test).

**Figure 5.**
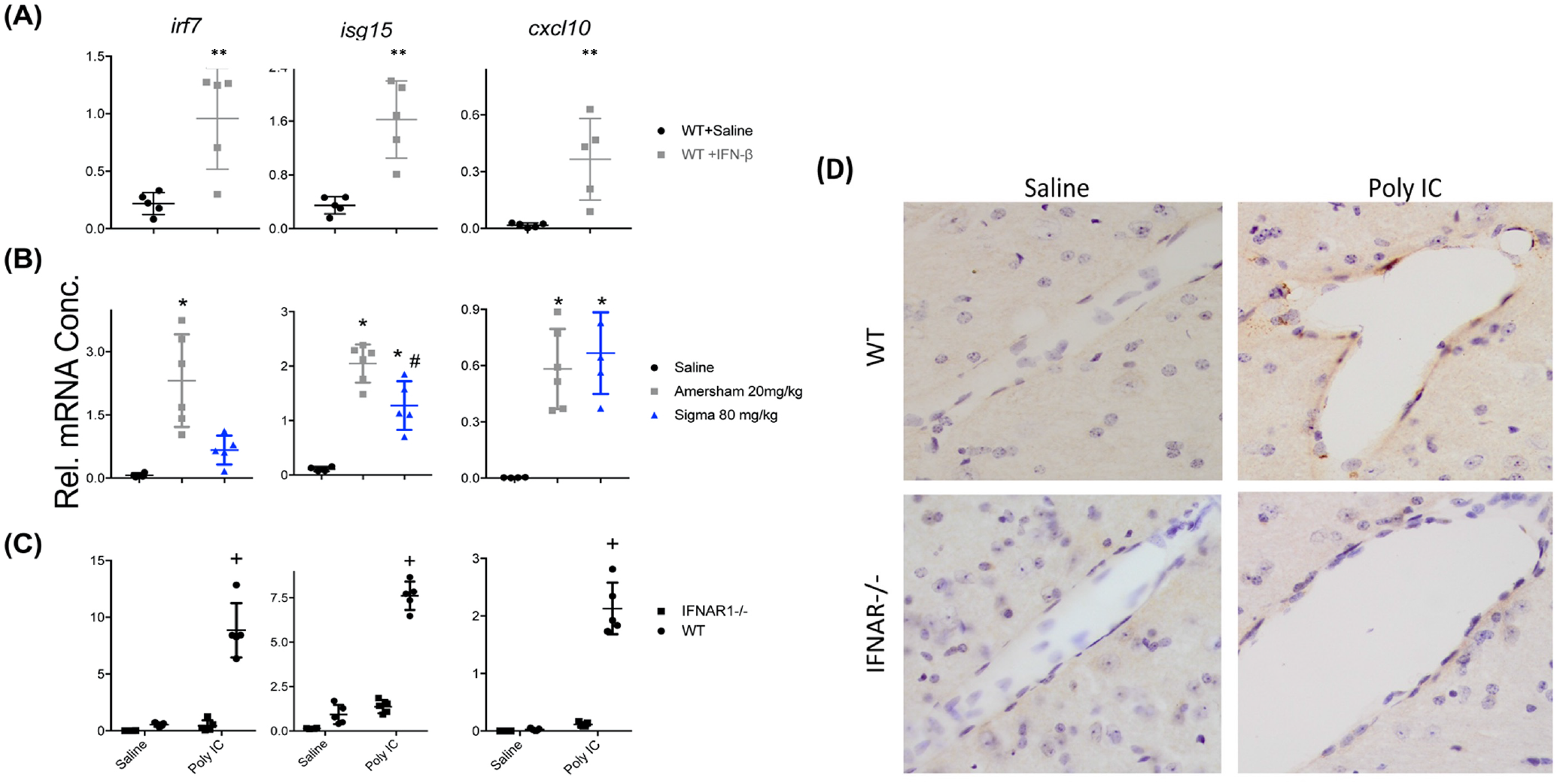
Relationship between circulating IFNβ and brain interferon-dependent responses. A) Hippocampal expression of interferon-dependent genes in adult female C57BL/6 mice treated with IFN-β (25,000 units i.p.; 3hr post injection) challenge compared to WT Saline treated (n=5 per group; unpaired Student’s t-test, ** p < 0.01). (B) Comparison of C57BL/6 mice challenged with saline, HMW (20 mg/kg) or LMW (80 mg/kg) poly I:C at 5hr post injection (n=4,6,5 for saline, HMW and LMW respectively; one-way ANOVA followed by Bonferroni post-hoc analyses: * indicates p < 0.05 when compared to saline and # indicates p < 0.05 when comparing both poly IC treatments). (C) Comparison of poly IC (HMW 20 mg/kg), 3 hrs post injection) in C57BL/6 mice and IFNAR1^-/-^ mice (all groups n=5 except WT saline n=4; + denotes significant difference between IFNAR1^-/-^ poly IC and WT poly IC and also between IFNAR1^-/-^ poly I:C and IFNAR1^-/-^ saline (p < 0.05) (D) Representative bright field photomicrographs (40X) of CXCL10 immunohistochemistry showing CXCL10-positive cells at the vascular endothelium, present only in WT poly I:C group.

Challenging normal C57 mice with HMW poly I:C (Figure 5b), it was clear once again that *Irf7* expression in the hippocampus was expressed to a much lower extent with LMW poly I:C (p<0.01). Another interferon-dependent gene, *Isg15*, was also expressed less robustly after LMW poly I:C treatment (p<0.01), but levels were still significantly elevated with respect to saline controls (p<0.01). The interferon-dependent gene *Cxcl10* showed somewhat variable transcription but was equally induced by LMW poly I:C (80 mg/kg) and HMW poly I:C (20 mg/kg) despite quite different levels of circulating IFNβ (Figure 4a). This may indicate that the high expression of other cytokines such as IL-6, TNF-α or IL-1β may act cooperatively to induce its transcription.

Challenging WT and IFNAR1^-/-^ mice with HMW poly I:C (Figure 5c), it was apparent that IFNAR1 was essential for the expression of *Irf7* in the hippocampus. Although expression of both *Isg15* and *Cxcl10* were both very much reduced in IFNAR1^-/-^ mice, some limited expression of these genes remained, supporting a role for induction by cytokines other than type I IFN. Nonetheless, at the protein level, using immunohistochemistry we show that CXCL10 is expressed at the brain endothelium in WT mice treated with HMW poly I:C but is absent in IFNAR1^-/-^ mice (Figure 5d).

### Effect of poly I:C on innate immune responses in young and aged mice

Now focusing on HMW due to its robust IFN-I responses, we examined the impact of age upon IFN-I and other cytokine responses. Pro-inflammatory cytokines were measured in blood plasma (IL-6, IFNβ, TNF-α and IL-1β) from animals treated with saline or poly I:C (at 3 or 21 months). Statistically significant differences in responses in of these groups were assessed by two-way ANOVA with Bonferroni post hoc analysis.

IL 1β was significantly induced (Figure. 6a), showing a main effect of treatment (F_1,19_ = 18.18, p= 0.0005) and of age (F_1,19_= 6.67, p = 0.0194) with a trend towards interaction between these factors (F_1,19_= 3.46, p=0.08). Bonferroni post-hoc comparison showed that poly I:C at 21 months induced significantly more IL-1β than the same challenge at 3 months (p=0.025). Poly I:C also robustly increased TNF-α (fig. 6a; main effect of treatment: F_1,19_ = 24.44, p<0.0001). However, young and aged groups showed similar increases and there was no effect of age and no interaction between treatment and age. Poly I:C had a marked effect on IL-6 secretion (Figure 6a; main effect of treatment F_1,19_ = 58.97, p<0.0001, and of age: F_1,19_ = 25.96, p<0.0001). Importantly, there was also an interaction between treatment and age, illustrating that poly I:C has greater impacts on plasma IL-6 in those with more advanced age (F_1,19_ = 22.84, p=0.0001). Age also significantly affected IFNβ responses to poly I:C in IFN β (fig. 6b). There was a significant main effect of treatment (F_1,19_ = 49.35, p<0.0001) and of age (F_1,19_ = 12.28, p= 0.0024). Again there was an Interaction between treatment and age showing exaggerated IFNβ responses with advanced age (F_1,19_=12.28, p=0.0024). Therefore there is evidence for more intense IL-1β, IL-6 and IFNβ responses to poly I:C in aged animals compared to young ones.

**Figure 6.**
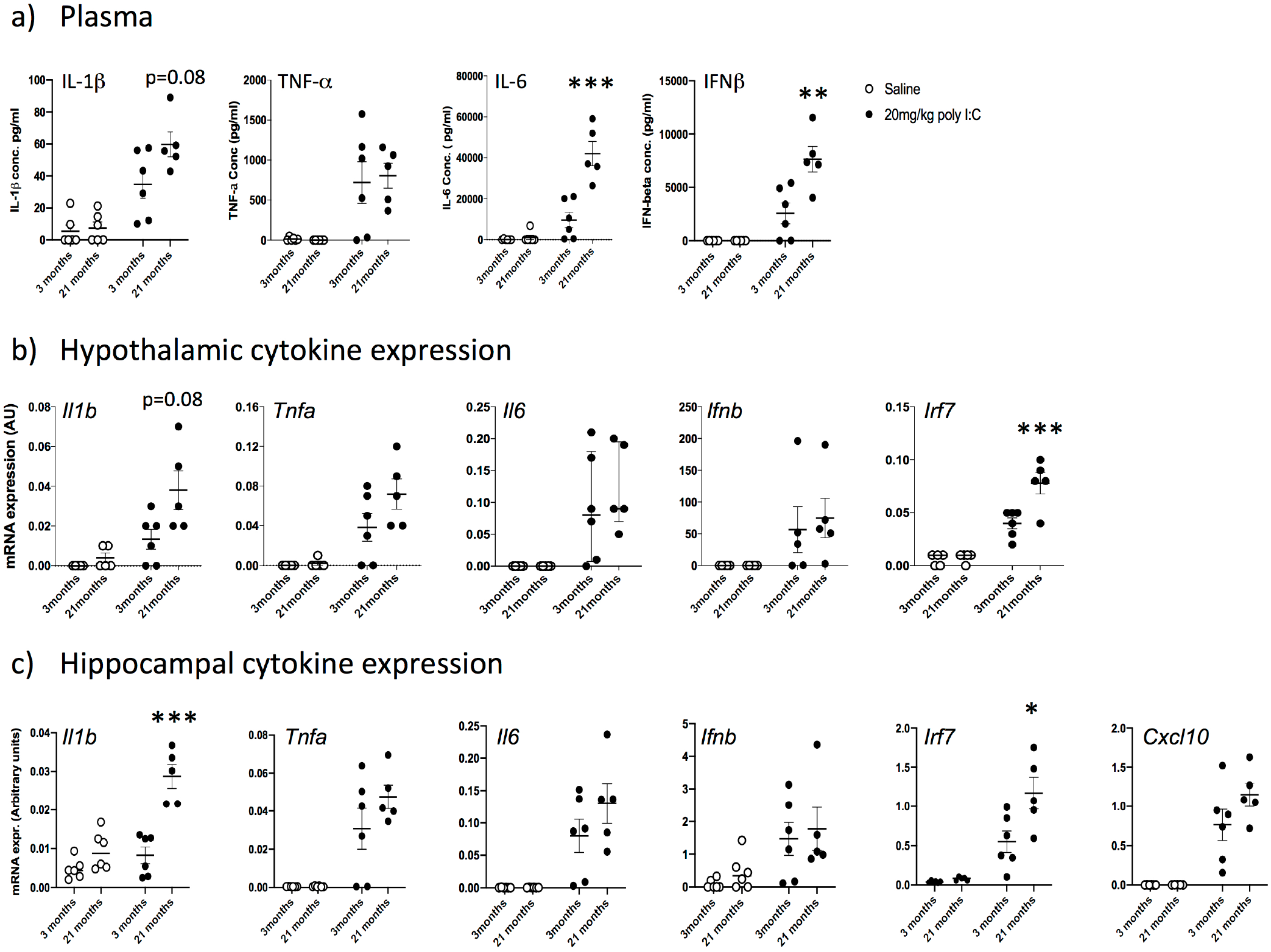
Impact of age on innate immune responses to poly I:C. a) Pro-inflammatory cytokines in plasma were measured by ELISA (IL-1β, TNF-α, IL-6 and IFNβ). Expression of cytokines in the hypothalamus (b) and hippocampus (c) was determined using quantitative PCR. All data were compared using two-way ANOVA followed by Bonferroni post-hoc tests. Poly I:C had a significant main effect on all cytokines in all areas. Significant interactions of treatment with age are indicated by *p<0.05, **p<0.01, ***p<0.001 and significant post-hoc differences between poly I:C in young and aged by # p<0.05 (n=6 for all groups except young+saline; n=5). All data are represented by mean+/− SEM.

Hypothalamic inflammatory transcripts (*Il1b, Tnfa, Il6, Ifnb, Irf7*) were measured in the same 3 month and 21 month old animals (Figure 6b). There were large effects of treatment for *Il1b, Tnfa* and *Il6* (F_1,17_ ≥18.18, p≤0.0005) while treatment also significantly affected *Ifnb* (F_1,17_=5.75, p=0.028). Among the cytokines, age was only a significant influence on *Il1b* (F_1,17_=6.67, p=0.0194), but the IFN-dependent gene *Irf7* also showed significant effects of age and treatment and, importantly, an interaction between these factors (F_1,17_=9.93, p=0.007). Bonferroni post-hoc analysis on all 5 transcripts showed that only *Il1b* (p=0.025) and *Irf7* (p=0.038) showed heightened responses in poly I:C at 21 months compared to 3 months.

Hippocampal inflammatory transcripts (*Il1b, Tnfa, Il6, Ifnb, Irf7*) were measured in the same animals (Figure 6c), with *Cxcl10* now added because of its reported impact on hippocampal-dependent cognitive function in recent studies (Blank *et al*., 2016). Transcription of *Il1b* was modest in young animals but marked in 21 month old mice. There was a significant interaction of treatment and age, with aged animals showing exaggerated *Il1b* expression (F_1,19_ = 29.19, p<0.0001). For *Tnfa*, (figure 6c) there was an effect of treatment and (F_1,19_ =38.31, p< 0.001) but levels were not different as a function of age and there was no interaction between age and treatment. Poly I:C produced quite variable increases in *Il6* in both young and aged animals (main effect of treatment F_1,19_ = 30.74) but there was no effect of age nor an interaction between age and treatment. *Ifnb*, expression in these samples was highly variable (qPCR assays for this single exon gene can be variable since they are prone to amplification of genomic DNA impurities in RNA) (Figure 6c). Poy I:C did induce *Ifnb* (main effect of treatment: F_1,19_ = 11.13, p=0.0035) but there was no effect of age and no interaction.

The *Irf7* response to poly I:C was exaggerated in aged mice with respect to young mice, (significant age x treatment interaction: F _1,18_ = 6.00, *p*=0.0248). Bonferroni post-hoc showed that poly I:C *Irf7* response at 21 months was significantly larger than that at 3 months (p<0.003). *Cxcl10* was also significantly elevated by poly I:C (main effect of treatment: F_1,18_ 54.85, P<0.0001) but there was no significant effect of age. There was a modest increase at 21 months with respect to 3 months but this was not quite significant (p=0.096). Therefore there is evidence of exaggerated *Il1b* response and exaggerated type I interferon action when aged mice are exposed to systemic double stranded RNA.

### Systemic poly I:C induces acute cognitive dysfunction selectively in aged animals

A separate batch of animals of 6 and 24 months of age were trained on an ‘escape from shallow water’ version of T-maze alternation, designed to test animals on working memory during concurrent sickness behavior (Murray *et al*., 2012). Mice were trained for >10 blocks of 10 trials in order to reach criterion performance of 80% correct responding and all animals, at baseline were capable of learning the alternation strategy to escape the maze regardless of age. Upon challenge with poly I:C (12 mg/kg i.p.) there was an acute disruption of working memory that occurred selectively in 24 month old animals challenged with poly I:C. The same challenge in young animals had no impact on cognitive function. There were main effects of age (F_1,68_=17.96) and poly I:C (F_1,68_=18.26, p<0.0001), but importantly there was an interaction of age and poly I:C (F_1,68_=4.733, p=0.033) illustrating that poly I:C has distinct effects on cognitive function in older animals.

## Discussion

In the current study we have shown that LMW dsRNA (such as that supplied by Sigma) induces much lower levels of pro-inflammatory cytokines IL-6 and TNF-α than HMW and induces negligible circulating IFNβ. Limited effects of LMW poly I:C on sickness behaviour can be overcome using significantly higher doses of poly I:C, but the refractory IFNβ responses persist. Time course studies indicate that brain IFN-I responses are directly dependent on circulating IFNβ and that brain *Ifnb* transcription is not, of itself, sufficient to trigger downstream anti-viral responses in the brain. IFNβ challenge is a strong driver of brain *Cxcl10* responses but both sickness and *Cxcl10* remain inducible in IFNAR1^-/-^ mice with higher LMW poly I:C dosing, consistent with recently described roles for CXCL10 in sickness behavior. Importantly IL-6, IL-1β and IFNβ or IFN-dependent responses were exaggerated in older animals, both in plasma and in brain and this was associated with significantly worse cognitive impairments in aged animals.

### Differential responses to HMW and LMW

We have shown widely different cytokine responses to Sigma and Amersham preparations of poly I:C and then used Invivogen poly I:C preparations, explicitly described, and verified here, as HMW or LMW, to show that Sigma preparations are LMW (<500 bp) and those of Amersham are HMW (1-6 kb). HMW poly I:C lead to spontaneous abortion in pregnant dams (table 1), which never occurred with LMW poly I:C, and this was recently reported elsewhere for HMW poly I:C (Mueller *et al*., 2019). This is likely to be caused by TNF-α since this was absent in LMW challenges at 20 mg/kg and TNF-α has been described to be a key mediator in inducing miscarriage in LPS and malaria models (Gendron *et al*., 1990; Poovassery *et al*., 2009). Two recent studies (Careaga *et al*., 2018; Mueller *et al*., 2019) have also suggested there is also significant variability between different Sigma preparations, showing variable amounts of higher MW dsRNA, which strongly predicted temperature and cytokine changes. Although HMW induced significantly greater cytokine levels (Mueller *et al*., 2019) neither study examined the prototypical anti-viral cytokine family, IFN-I. The data from all 3 studies show that cytokine responses require characterisation with each individual poly I:C preparation but the current data indicate that IFN-I responses are unlikely to have been prevalent in the majority of published studies of maternal immune activation that have been performed to date, since most of those studies use Sigma poly I:C preparation. IFN-I responses are important contributors to the sickness behaviour response in poly I:C treated animals (Murray *et al*., 2015; Blank *et al*., 2016) and also facilitate the IL-6 response. That is, while IL-6 appears to contribute to the hypolocomotion and reduction in species-typical behaviours associated with poly I:C induced sickness, IL-6 production is itself dependent on baseline IFN-I levels (Matsumoto & Seya, 2008b; Murray *et al*., 2015). IL-6 is an important mediator of poly I:C- and virally-induced behavioural changes in the offspring of maternal immune activation models (Matsumoto & Seya, 2008b; Murray *et al*., 2015) so it is apparent that levels of IFN-I and IL-6 are intimately linked and models purporting to mimic viral infection must be cognisant of this deficiency as well as the variability inherent in LMW preparations.

### Different cytokine responses: different receptors and different locations?

The fundamentally different cytokine responses observed here, and in recent studies, showing quantitative (Careaga *et al*., 2018; Mueller *et al*., 2019) and qualitative (IFN-I: current study) differences, suggest that distinct receptors are being engaged by these different preparations. Synthetic dsRNA, poly I:C, activates the immune system by signaling through TLR3, RIG-I and MDA5. TLR3 is found on the endosomal lumen of innate immune cells. Cell surface TLR3 expression is also seen on fibroblast and epithelial cells but successful poly I:C-TLR3 interactions are dependent on an acidic pH within the endosome (de Bouteiller *et al*., 2005; Matsumoto & Seya, 2008a; Gürtler & Bowie, 2013). Macrophages and, in particular, plasmacytoid dendritic cells can take up extracellularly applied poly I:C, reflecting their essential ‘sentinel’ response to viral infection of the host, while other cell types require transfection of RNA in order to access cytoplasmic dsRNA sensors and induce IFNβ responses. Given their primary role in recognising viral infection, it is significant that pDCs and macrophages do not require transfection (Zhou *et al*., 2013).

Both RIG-I and MDA5 are cytoplasmic and thus dsRNA must reach the cytoplasm to facilitate sensing by these receptors. Cytoplasmic delivery (with transection agents such as polyethylenimine (PEI) derivates) was required to trigger MDA5 responses but not TLR3 responses (Kato et al., 2008). *In vitro*, naked dsRNA requires TLR3 while liposome encapsulated poly I:C does not, since it has access to the cytoplasm (Dauletbaev *et al*., 2015). Most neuroscience studies with poly I:C do not use transfection agents. Prinz and colleagues (Blank *et al*., 2016) used jetPEI to ‘transfect’ multiple cell types, *in vivo*, with poly I:C and showed a requirement for mitochondrial antiviral signalling protein (MAVS) and the IFNAR1 receptor to induce sickness behaviour and cognitive effects of poly I:C and VSV infection. Other authors using 10 mg/Kg poly I:C i.p., without transfection (similar to current study), still found MAVS to be essential in inducing IFN-I and the full IL-6 response (Sun *et al*., 2006). Thus, cytosolic sensing of dsRNA is essential for full responses and MAVS is important whether transfection of poly I:C has been performed (Blank *et al*., 2016) or not (Sun *et al*., 2006). This implicates either RIG-I or MDA5 in the current study.

The length of dsRNA induces different IFN-I and cytokine responses to virally-generated dsRNA (DeWitte-Orr *et al*., 2009) and it is now clear that MDA5 binds long dsRNA while RIG-I preferentially recognises LMW: while extremely similar in structure, RIG-I binds to the blunt end or 5’ triphosphate (5’ ppp) end of dsRNA while MDA-5 alternatively recognises the duplex structure of dsRNA rather than any particular base pair sequence and is particularly sensitive to long strands (Kato *et al*., 2008). Despite their different dsRNA length preferences, MDA-5 and RIG-I signalling converge on MAVS signalling, recruiting dsRNA to the mitochondria where it acts, via TBK-1 and IKK-i, to induce NF-κB and IRF3 (Gürtler and Bowie, 2013). Though RIG-I and MDA5 can both induce IFNβ (Dauletbaev *et al*., 2015; Palchetti *et al*., 2015) HMW appears necessary for IFNβ (Zhou *et al*., 2013). LMW cannot induce IFNβ expression in various cell types without transfection and the lower efficiency of cytoplasmic exposure to LMW dsRNA (Zhou *et al*., 2013) may be sufficient to effectively allow *Ifnb* induction to occur only at trivial levels with LMW poly I:C in the current study. However in TLR3 replete cells, MAVS is not necessary for initial pDC responses to poly IC (Sun *et al*., 2006) and TLR3 activation and IFNβ production by HMW poly I:C is higher than that with LMW *in vitro* (Zhou *et al*., 2013). Based on the above findings, and the data suggesting TLR3 also contributes to influenza-induced sickness behaviour (Majde *et al*., 2009), its likely that TLR3 has a role in facilitating organismal responses to extracellularly available dsRNA, but that MDA5 plays the major role in cytokine and IFNβ induction. This may occur upon sensing poly I:C distributed to the cytoplasm by extracellular vesicles (Frank-Bertoncelj *et al*., 2018) or other routes.

### Brain responses, Sickness behaviour, IFN-I and CXCL10

Systemic IFNβ responses and brain expression of IFN-dependent genes were minimal with LMW poly I:C despite clear evidence of transcription of *Ifnb* and other cytokine transcripts in brain homogenates, even at lower LMW doses (figure 2). Since these animals had very low levels of circulating cytokines it seems reasonable to conclude that these brain transcripts represent a direct endothelial and epithelial response to 20 mg/kg poly I:C appearing in the circulation. Despite clear *Ifnb* transcription arising from both LMW and HMW treatment (becoming even higher than with HMW poly I:C at high concentrations of LMW), LMW did not robustly induce *Irf7*, which is a definitive indicator of local response to active IFNβ (Gurtler & Bowie, 2013). This indicates that the brain IFN-dependent response (*Irf7*) is in response to circulating IFN-I, as validated in figure 5 and consistent with the observation of widespread brain interferon-dependent gene expression in response to IFNα (Wang *et al*., 2008). The brain *Irf7* response to HMW poly I:C was completely ablated in IFNAR1^-/-^ mice, as was *Cxcl10*, but *Cxcl10* expression was robustly induced by LMW poly I:C when this was applied at high doses, even when *Irf7* was minimally expressed. This reflects the finding that *Cxcl10* expression can also be induced directly by poly I:C, regulated by NFκB (Brownell *et al*., 2014). This failure of transcribed *Ifnb* to induce IFN-dependent responses may be because type I IFN signaling in the absence of other cues activates the expression of IFN-inducible genes that make cells more sensitive to further virus-induced signals (Stetson & Medzhitov, 2006). That could mean that dsRNA recognition triggers *Ifnb* transcription in cells that are unable to complete the synthesis of IFNβ to induce downstream programs without additional recognition of circulating IFNβ. There is some evidence of post-transcriptional regulation of IFN-I (Khabar & Young, 2007). There is also evidence that the brain responds differently to poly I:C compared to peripheral tissues, with increasing *Ifnb* expression in MyD88^-/-^ after icv poly I:C (Zhu *et al*., 2016) but this does not offer a satisfactory explanation for the failure to express *Irf7* downstream of *Ifnb* transcription after LMW poly I:C challenge.

Although we found that even at high doses LMW poly I:C did not produce significant IFNβ, these animals nonetheless became significantly more sick than the IFNβ-inducing HMW poly I:C at 20 mg/kg. That implies that, although IFNAR1 may be central to sickness behaviour responses to systemic poly I:C (Murray *et al*., 2015; Blank *et al*., 2016), high levels of other cytokines are sufficient to drive the sickness response in the absence of circulating IFNβ. Although the study of Blank et al shows that IFN-dependent CXCL10 drives the sickness behaviour response, we have previously shown that elements of the sickness response that are IFNAR1-dependent can be recapitulated, in IFNAR1^-/-^ mice, by supplementation with IL-6 (Murray et al., 2015). Here, treatment with high dose of LMW poly I:C fails to produce circulating IFNβ but does induce very high levels of circulating IL-6 (figure 3,4) and also robust brain induction of *Cxcl10* despite the lack of IFNβ (Figure 5). The data can thus be reconciled with prior data demonstrating the role of IFNβ and CXCL10 in poly I:C-induced sickness behaviour. Further studies are required to determine whether the induction of CXCL10 in the absence of IFNβ or IL-6 is sufficient for the production of sickness behaviour.

### Exaggerated responses to dsRNA with age and relevance to viral infection in the elderly

There is significant interest in determining whether aged individuals respond differently to equivalent exposure to dsRNA, as might occur with viral infections encountered by the older population. The current data demonstrate exaggerated IFN-I and cytokine responses to poly I:C in aged mice (Figure 6). Both IL-6 and IFNβ show exaggerated induction in the blood while *Il1b* and *Irf7* (indicating IFN-I action) showed exaggerated induction in the hippocampus. The molecular basis for this exaggerated cytokine response has not been elucidated here, but there is evidence that low level IFN-I expression facilitates more robust IFN-I and IL-6 responses to subsequent challenge (Gough *et al*., 2012) and there is existing evidence for low-grade IFN-I expression in aged mice (Baruch *et al*., 2014). Moreover, microglia are known to be primed to produced exaggerated cytokine responses in aged mice (Godbout *et al*., 2005).

These data are important in a general sense since many viruses shed dsRNA during infection and the innate immune response occurring during viral infection may be a key driver of acute cognitive deficits in infected individuals. The current data indicate that, at least with respect to dynamic cognitive processes like working memory, aged animals are significantly more susceptible to these changes upon exposure to dsRNA (Figure 7). These cognitive data, have particular resonance in the current coronavirus SARS-CoV-2 pandemic, in which CNS symptoms including confusion and delirium have been widely observed. In the largest report on hospitalised cases of SARS-CoV-2 in Europe, the ISARIC consortium report that confusion was the most 5^th^ most common symptom on admission, representing 25% of all patients (Docherty *et al*., 2020) (Docherty *et al*., 2020; Varatharaj *et al*., 2020). It is known that delirium is more prevalent in older age (Cole, 2004) and that older SARS-CoV-2 patients often present with delirium in the absence of even the characteristic COVID19 symptoms (Kennedy *et al*., 2020; Poloni *et al*., 2020). There is, quite rightly, significant attention on the evidence that SARS-CoV-2 can enter the brain (Meinhardt *et al*., 2020) and whether this contributes to the frequent neuropsychiatric features observed, but the current data show that even in the absence viable virus, peripheral dsRNA alone is sufficient to trigger disruption of dynamic cognitive functions relying on attention and working memory, selectively in older individuals.

**Figure 7.**
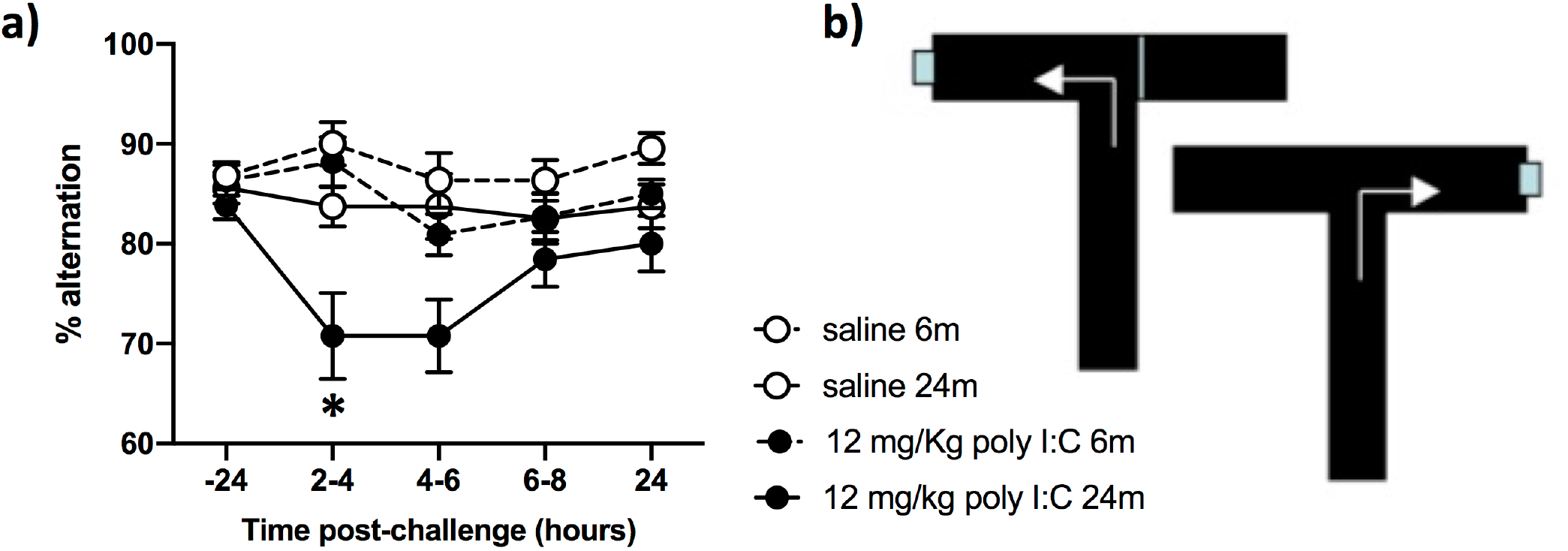
Selective effects of poly I:C on cognitive function in older animals. a) Animals were trained for 10 days x 10 trials per day (on the escape-from-water T-maze shown in b) and then tested at baseline (−24 h) before challenge with poly I:C at 12 mg/kg i.p. or sterile saline and testing for 3 blocks of 5 trials between 2-4h, 4-6h, 6-8h and 24h. Data are presented as mean±SEM (Saline 6m n=22, saline 24m n=15, poly I:C 6 m n=22, poly I:C n=13). Data were analysed by 3 way ANOVA and * denotes an interaction of age and poly I:C.

Double stranded RNA is a relevant immune stimulus with respect to SARS-CoV-2, an enveloped positive sense ssRNA virus (like flaviviruses, retroviruses). Once inside the cytoplasm +sense RNA is read by ribosomes to make viral proteins and acting as a template to form an intermediate hybrid state with negative sense RNA to facilitate making multiple copies of the original +ssRNA. Thus significant dsRNA is present for a period, which allows the activation of dsRNA sensors such MDA5 (Kato *et al*., 2008) and with a genome of approximately 30 kb (Bar-On *et al*., 2020) long RNA (i.e. HMW) duplexes are likely to occur. Nothwithstanding evidence that coronaviruses can form double membrane vesicles that sequester dsRNA and limit their engagement of PRRs (Blanco-Melo *et al*., 2020; Vabret *et al*., 2020), RNAemia was detected in 78.6% of SARS-CoV-2 patients who required hospital admission (Hogan *et al*., 2020). There is a type I interferon response in SARS-CoV-2 infections in humans. Not all studies concur (Hadjadj *et al*., 2020), but recent scRNAseq evidence suggests that IFN-I, along with IL-1- and TNF-dependent pathways may be particularly important in severe infection (Lee *et al*., 2020).

Whether there are heightened IFN-I responses in aged individuals exposed to viral infection more generally requires investigation, but it remains striking that aged animals show exaggerated pro-inflammatory and IFN-I responses and also show exaggerated vulnerability to acute cognitive deficits relevant to delirium. This selective disruption of working memory in aged animals, and its relevance to viral infection-induced delirium requires investigation. The increased risk of delirium in aged patients is true for many infections, including influenza, but information on specific viral infections is scant and this requires investigation. There is prior evidence for poly I:C-induced memory impairments. Poly I:C affected contextual fear conditioning memory consolidation in normal animals (Weintraub *et al*., 2014) and more detailed studies show motor learning deficits and spine remodelling that was dependent on TNF-α and peripheral monocytes (Garre *et al*., 2017), most likely with LMW poly I:C. While aged mice showed slightly increased sickness behaviour responses to poly I:C compared to younger animals (McLinden *et al*., 2012) age-dependent impacts of dsRNA on cognitive function such as those shown here, have not previously been described.

Arguing for a cytokine-mediated basis for the acute cognitive changes seen here, we have previously shown direct causative roles of IL-1β (Skelly *et al*., 2019) and TNF-α (Hennessy *et al*., 2017) in similar deficits and there are also now a number of studies indicating that IFNβ can also have direct or indirect impacts on neuronal physiology and cognition (Costello *et al*., 2011; Baruch *et al*., 2014; Blank *et al*., 2016). Given the current trials of IFNα or β in SARS-CoV2 patients, it is very important to elucidate the cognitive effects of IFN-I in aged individuals and patients with cognitive impairment at baseline.

We posit that at least some of the very commonly observed CNS disturbances associated with SARS-CoV-2 can be understood in the context of well-described systemic inflammation causing acute cognitive dysfunction when superimposed on the vulnerable brain. The contribution of SARS-CoV-2-induced systemic inflammation is also important to examine in the context of new brain injury and acceleration of dementia and cognitive aging. We have shown that repeated poly I:C treatments produce acute episodes of dysfunction but also accelerate neurological decline (Field *et al*., 2010) and this mirrors the reported impact of delirium on subsequent dementia diagnoses and rate of decline (Fong *et al*., 2009; Davis *et al*., 2012). The recent description of increased markers of brain injury in SARS-CoV-2 patients (Kanberg *et al*., 2020) is consistent with the view that this infection produces new brain injury that may accelerate cognitive decline. Supporting a possible role for IFN-I in such disease exacerbation we (Nazmi *et al*., 2019) and others (Taylor *et al*., 2014; Abdullah *et al*., 2018) have shown that brain IFNAR1 contributes to progression and severity of neurodegenerative disease. It is important to add that while inflammatory cytokines can directly impact on brain function, hypercoagulation, which is triggered by inflammation-induced microvascular damage, is likely an important determinant of more severe outcomes in SARS-CoV-2 patients (Liao *et al*., 2020) and post-mortem studies showing these features are beginning to emerge (Nath *et al*., 2020). Hypercoagulation can also be induced by TLR3 stimulation (Shibamiya *et al*., 2009). Mechanisms by which systemic viral infections may contribute to neurodegeneration require investigation.

## Conclusion

HMW poly I:C has substantially more robust effects on systemic and brain inflammation than LMW poly I:C in C57BL6 mice. In addition, LMW poly I:C fails to induce a robust type I IFN response, the prototypical cytokine system of the antI-viral response. The data showing that circulating IFNβ is the major determinant of brain IFN-I responses, that this response is exaggerated in aged mice and that working memory disruption is exaggerated in aged mice treated with poly I:C has implications for understanding acute cognitive effects of viral infections in older individuals and raise questions about the contribution of systemic viral infections to neurodegenerative changes.

## Acknowledgements

Dr. Niamh McGarry was supported by a Ph.D. studentship under the PRTLI programme “Molecular and Cellular Mechanisms underlying Inflammatory Processes”. Additional funding for the work was generously provided by the Wellcome Trust as part of a SRF (090907) to CC and by the NIH (R01AG0506236).

## References

Abdullah, A., Zhang, M., Frugier, T., Bedoui, S., Taylor, J.M. & Crack, P.J. (2018) STING-mediated type-I interferons contribute to the neuroinflammatory process and detrimental effects following traumatic brain injury. J Neuroinflammation, 15, 323.

Alexopoulou, L., Holt, A.C., Medzhitov, R. & Flavell, R.A. (2001) Recognition of double-stranded RNA and activation of NF-kappaB by Toll-like receptor 3. Nature, 413, 732–738.

Bar-On, Y.M., Flamholz, A., Phillips, R. & Milo, R. (2020) SARS-CoV-2 (COVID-19) by the numbers. Elife, 9.

Baruch, K., Deczkowska, A., David, E., Castellano, J.M., Miller, O., Kertser, A., Berkutzki, T., Barnett-Itzhaki, Z., Bezalel, D., Wyss-Coray, T., Amit, I. & Schwartz, M. (2014) Aging-induced type I interferon response at the choroid plexus negatively affects brain function. Science, 346, 89–93.

Blanco-Melo, D., Nilsson-Payant, B.E., Liu, W.C., Uhl, S., Hoagland, D., Moller, R., Jordan, T.X., Oishi, K., Panis, M., Sachs, D., Wang, T.T., Schwartz, R.E., Lim, J.K., Albrecht, R.A. & tenOever, B.R. (2020) Imbalanced Host Response to SARS-CoV-2 Drives Development of COVID-19. Cell, 181, 1036–1045 e1039.

Blank, T., Detje, C.N., Spiess, A., Hagemeyer, N., Brendecke, S.M., Wolfart, J., Staszewski, O., Zoller, T., Papageorgiou, I., Schneider, J., Paricio-Montesinos, R., Eisel, U.L., Manahan-Vaughan, D., Jansen, S., Lienenklaus, S., Lu, B., Imai, Y., Muller, M., Goelz, S.E., Baker, D.P., Schwaninger, M., Kann, O., Heikenwalder, M., Kalinke, U. & Prinz, M. (2016) Brain Endothelial- and Epithelial-Specific Interferon Receptor Chain 1 Drives Virus-Induced Sickness Behavior and Cognitive Impairment. Immunity, 44, 901–912.

Brownell, J., Bruckner, J., Wagoner, J., Thomas, E., Loo, Y.M., Gale, M., Jr., Liang, T.J. & Polyak, S.J. (2014) Direct, interferon-independent activation of the CXCL10 promoter by NF-kappaB and interferon regulatory factor 3 during hepatitis C virus infection. J Virol, 88, 1582–1590.

Careaga, M., Taylor, S.L., Chang, C., Chiang, A., Ku, K.M., Berman, R.F., Van de Water, J.A. & Bauman, M.D. (2018) Variability in PolyIC induced immune response: Implications for preclinical maternal immune activation models. J Neuroimmunol, 323, 87–93.

Cole, M.G. (2004) Delirium in elderly patients. Am J Geriatr Psychiatry, 12, 7–21.

Costello, D.A., Lyons, A., Denieffe, S., Browne, T.C., Cox, F.F. & Lynch, M.A. (2011) Long term potentiation is impaired in membrane glycoprotein CD200-deficient mice: a role for Toll-like receptor activation. J Biol Chem, 286, 34722–34732.

Crum, W., Sawiak, S., Chege, W., Cooper, J., Williams, S. & Vernon, A. (2017) Evolution of structural abnormalities in the rat brain following in utero exposure to maternal immune activation: A longitudinal in vivo MRI study. Brain, Behavior and Immunity, 63, 50–59.

Cunningham, C., Campion, S., Teeling, J., Felton, L. & Perry, V.H. (2007) The sickness behaviour and CNS inflammatory mediator profile induced by systemic challenge of mice with synthetic double-stranded RNA (poly I:C). Brain Behav Immun, 21, 490–502.

Dauletbaev, N., Cammisano, M., Herscovitch, K. & Lands, L.C. (2015) Stimulation of the RIG-I/MAVS Pathway by Polyinosinic:Polycytidylic Acid Upregulates IFN-beta in Airway Epithelial Cells with Minimal Costimulation of IL-8. J Immunol, 195, 2829–2841.

Davis, D.H., Muniz Terrera, G., Keage, H., Rahkonen, T., Oinas, M., Matthews, F.E., Cunningham, C., Polvikoski, T., Sulkava, R., MacLullich, A.M. & Brayne, C. (2012) Delirium is a strong risk factor for dementia in the oldest-old: a population-based cohort study. Brain, 135, 2809–2816.

de Bouteiller, O., Merck, E., Hasan, U.A., Hubac, S., Benguigui, B., Trinchieri, G., Bates, E.E. & Caux, C. (2005) Recognition of double-stranded RNA by human toll-like receptor 3 and downstream receptor signaling requires multimerization and an acidic pH. Journal of Biological Chemistry, 280, 38133–38145.

DeWitte-Orr, S.J., Mehta, D.R., Collins, S.E., Suthar, M.S., Gale, M., Jr. & Mossman, K.L. (2009) Long double-stranded RNA induces an antiviral response independent of IFN regulatory factor 3, IFN-beta promoter stimulator 1, and IFN. J Immunol, 183, 6545–6553.

Docherty, A.B., Harrison, E.M., Green, C.A., Hardwick, H.E., Pius, R., Norman, L., Holden, K.A., Read, J.M., Dondelinger, F., Carson, G., Merson, L., Lee, J., Plotkin, D., Sigfrid, L., Halpin, S., Jackson, C., Gamble, C., Horby, P.W., Nguyen-Van-Tam, J.S., Ho, A., Russell, C.D., Dunning, J., Openshaw, P.J., Baillie, J.K., Semple, M.G. & investigators, I.C. (2020) Features of 20 133 UK patients in hospital with covid-19 using the ISARIC WHO Clinical Characterisation Protocol: prospective observational cohort study. BMJ, 369, m1985.

Eyre, D.W. & Lumley, S.F. & O’Donnell, D. & Campbell, M. & Sims, E. & Lawson, E. & Warren, F. & James, T. & Cox, S. & Howarth, A. & Doherty, G. & Hatch, S.B. & Kavanagh, J. & Chau, K.K. & Fowler, P.W. & Swann, J. & Volk, D. & Yang-Turner, F. & Stoesser, N. & Matthews, P.C. & Dudareva, M. & Davies, T. & Shaw, R.H. & Peto, L. & Downs, L.O. & Vogt, A. & Amini, A. & Young, B.C. & Drennan, P.G. & Mentzer, A.J. & Skelly, D.T. & Karpe, F. & Neville, M.J. & Andersson, M. & Brent, A.J. & Jones, N. & Martins Ferreira, L. & Christott, T. & Marsden, B.D. & Hoosdally, S. & Cornall, R. & Crook, D.W. & Stuart, D.I. & Screaton, G. & Oxford University Hospitals Staff Testing, G. & Watson, A.J. & Taylor, A. & Chetwynd, A. & Grassam-Rowe, A. & Mighiu, A.S. & Livingstone, A. & Killen, A. & Rigler, C. & Harries, C. & East, C. & Lee, C. & Mason, C.J. & Holland, C. & Thompson, C. & Hennesey, C. & Savva, C. & Kim, D.S. & Harris, E.W. & McGivern, E.J. & Qian, E. & Rothwell, E. & Back, F. & Kelly, G. & Watson, G. & Howgego, G. & Chase, H. & Danbury, H. & Laurenson-Schafer, H. & Ward, H.L. & Hendron, H. & Vorley, I.C. & Tol, I. & Gunnell, J. & Ward, J.L. & Drake, J. & Wilson, J.D. & Morton, J. & Dequaire, J. & O’Byrne, K. & Motohashi, K. & Harper, K. & Ravi, K. & Millar, L.J. & Peck, L.J. & Oliver, M. & English, M.R. & Kumarendran, M. & Wedlich, M. & Ambler, O. & Deal, O.T. & Sweeney, O. & Cowie, P. & Naude, R.T.W. & Young, R. & Freer, R. & Scott, S. & Sussmes, S. & Peters, S. & Pattenden, S. & Waite, S. & Johnson, S.A. & Kourdov, S. & Santos-Paulo, S. & Dimitrov, S. & Kerneis, S. & Ahmed-Firani, T. & King, T.B. & Ritter, T.G. & Foord, T.H. & De Toledo, Z. & Christie, T. & Gergely, B. & Axten, D. & Simons, E.J. & Nevard, H. & Philips, J. & Szczurkowska, J. & Patel, K. & Smit, K. & Warren, L. & Morgan, L. & Smith, L. & Robles, M. & McKnight, M. & Luciw, M. & Gates, M. & Sande, N. & Turford, R. & Ray, R. & Rughani, S. & Mitchell, T. & Bellinger, T. & Wharton, V. & Justice, A. & Jesuthasan, G. & Wareing, S. & Huda Mohamad Fadzillah, N. & Cann, K. & Kirton, R. & Sutton, C. & Salvagno, C. & G, D.A. & Pill, G. & Butcher, L. & Rylance-Knight, L. & Tabirao, M. & Moroney, R. & Wright, S. & Peto, T.E. & Holthof, B. & O’Donnell, A.M. & Ebner, D. & Conlon, C.P. & Jeffery, K. & Walker, T.M. (2020) Differential occupational risks to healthcare workers from SARS-CoV-2 observed during a prospective observational study. Elife, 9.

Field, R., Campion, S., Warren, C., Murray, C. & Cunningham, C. (2010) Systemic challenge with the TLR3 agonist poly I:C induces amplified IFNalpha/beta and IL-1beta responses in the diseased brain and exacerbates chronic neurodegeneration. Brain Behav Immun, 24, 996–1007.

Fong, T.G., Jones, R.N., Shi, P., Marcantonio, E.R., Yap, L., Rudolph, J.L., Yang, F.M., Kiely, D.K. & Inouye, S.K. (2009) Delirium accelerates cognitive decline in Alzheimer disease. Neurology, 72, 1570–1575.

Fortier, M.E., Kent, S., Ashdown, H., Poole, S., Boksa, P. & Luheshi, G.N. (2004) The viral mimic, polyinosinic:polycytidylic acid, induces fever in rats via an interleukin-1-dependent mechanism. Am J Physiol Regul Integr Comp Physiol, 287, R759–766.

Frank-Bertoncelj, M., Pisetsky, D.S., Kolling, C., Michel, B.A., Gay, R.E., Jungel, A. & Gay, S. (2018) TLR3 Ligand Poly(I:C) Exerts Distinct Actions in Synovial Fibroblasts When Delivered by Extracellular Vesicles. Front Immunol, 9, 28.

Garay, P.A., Hsiao, E.Y., Patterson, P.H. & McAllister, A.K. (2013) Maternal immune activation causes age- and region-specific changes in brain cytokines in offspring throughout development. Brain Behav Immun, 31, 54–68.

Garre, J.M., Silva, H.M., Lafaille, J.J. & Yang, G. (2017) CX3CR1(+) monocytes modulate learning and learning-dependent dendritic spine remodeling via TNF-alpha. Nat Med, 23, 714–722.

Gendron, R.L., Nestel, F.P., Lapp, W.S. & Baines, M.G. (1990) Lipopolysaccharide-induced fetal resorption in mice is associated with the intrauterine production of tumour necrosis factor-alpha. J Reprod Fertil, 90, 395–402.

Godbout, J.P., Chen, J., Abraham, J., Richwine, A.F., Berg, B.M., Kelley, K.W. & Johnson, R.W. (2005) Exaggerated neuroinflammation and sickness behavior in aged mice following activation of the peripheral innate immune system. Faseb J, 19, 1329–1331.

Gough, D.J., Messina, N.L., Clarke, C.J., Johnstone, R.W. & Levy, D.E. (2012) Constitutive type I interferon modulates homeostatic balance through tonic signaling. Immunity, 36, 166–174.

Gurtler, C. & Bowie, A.G. (2013) Innate immune detection of microbial nucleic acids. Trends in microbiology, 21, 413–420.

Gürtler, C. & Bowie, A.G. (2013) Innate immune detection of microbial nucleic acids. Trends in microbiology, 21, 413–420.

Hadjadj, J., Yatim, N., Barnabei, L., Corneau, A., Boussier, J., Smith, N., Pere, H., Charbit, B., Bondet, V., Chenevier-Gobeaux, C., Breillat, P., Carlier, N., Gauzit, R., Morbieu, C., Pene, F., Marin, N., Roche, N., Szwebel, T.A., Merkling, S.H., Treluyer, J.M., Veyer, D., Mouthon, L., Blanc, C., Tharaux, P.L., Rozenberg, F., Fischer, A., Duffy, D., Rieux-Laucat, F., Kerneis, S. & Terrier, B. (2020) Impaired type I interferon activity and inflammatory responses in severe COVID-19 patients. Science, 369, 718–724.

Hennessy, E., Gormley, S., Lopez-Rodriguez, A.B., Murray, C. & Cunningham, C. (2017) Systemic TNF-alpha produces acute cognitive dysfunction and exaggerated sickness behavior when superimposed upon progressive neurodegeneration. Brain, behavior, and immunity, 59, 233–244.

Hogan, C.A., Stevens, B.A., Sahoo, M.K., Huang, C., Garamani, N., Gombar, S., Yamamoto, F., Murugesan, K., Kurzer, J., Zehnder, J. & Pinsky, B.A. (2020) High Frequency of SARS-CoV-2 RNAemia and Association With Severe Disease. Clin Infect Dis.

Kanberg, N., Ashton, N.J., Andersson, L.M., Yilmaz, A., Lindh, M., Nilsson, S., Price, R.W., Blennow, K., Zetterberg, H. & Gisslen, M. (2020) Neurochemical evidence of astrocytic and neuronal injury commonly found in COVID-19. Neurology, 95, e1754–e1759.

Kato, H., Takeuchi, O., Mikamo-Satoh, E., Hirai, R., Kawai, T., Matsushita, K., Hiiragi, A., Dermody, T.S., Fujita, T. & Akira, S. (2008) Length-dependent recognition of double-stranded ribonucleic acids by retinoic acid-inducible gene-I and melanoma differentiation-associated gene 5. J Exp Med, 205, 1601–1610.

Kennedy, M., Helfand, B.K.I., Gou, R.Y., Gartaganis, S.L., Webb, M., Moccia, J.M., Bruursema, S.N., Dokic, B., McCulloch, B., Ring, H., Margolin, J.D., Zhang, E., Anderson, R., Babine, R.L., Hshieh, T., Wong, A.H., Taylor, R.A., Davenport, K., Teresi, B., Fong, T.G. & Inouye, S.K. (2020) Delirium in Older Patients With COVID-19 Presenting to the Emergency Department. JAMA Netw Open, 3, e2029540.

Khabar, K.S. & Young, H.A. (2007) Post-transcriptional control of the interferon system. Biochimie, 89, 761–769.

Konat, G.W., Borysiewicz, E., Fil, D. & James, I. (2009) Peripheral challenge with double-stranded RNA elicits global up-regulation of cytokine gene expression in the brain. J Neurosci Res, 87, 1381–1388.

Lee, J.S., Park, S., Jeong, H.W., Ahn, J.Y., Choi, S.J., Lee, H., Choi, B., Nam, S.K., Sa, M., Kwon, J.S., Jeong, S.J., Lee, H.K., Park, S.H., Park, S.H., Choi, J.Y., Kim, S.H., Jung, I. & Shin, E.C. (2020) Immunophenotyping of COVID-19 and influenza highlights the role of type I interferons in development of severe COVID-19. Sci Immunol, 5.

Liao, D., Zhou, F., Luo, L., Xu, M., Wang, H., Xia, J., Gao, Y., Cai, L., Wang, Z., Yin, P., Wang, Y., Tang, L., Deng, J., Mei, H. & Hu, Y. (2020) Haematological characteristics and risk factors in the classification and prognosis evaluation of COVID-19: a retrospective cohort study. Lancet Haematol, 7, e671–e678.

Majde, J.A., Kapas, L., Bohnet, S.G., De, A. & Krueger, J.M. (2009) Attenuation of the influenza virus sickness behavior in mice deficient in Toll-like receptor 3. Brain Behav Immun.

Matsumoto, M. & Seya, T. (2008a) TLR3: interferon induction by double-stranded RNA including poly (I: C). Advanced drug delivery reviews, 60, 805–812.

Matsumoto, M. & Seya, T. (2008b) TLR3: interferon induction by double-stranded RNA including poly(I:C). Adv Drug Deliv Rev, 60, 805–812.

McLinden, K.A., Kranjac, D., Deodati, L.E., Kahn, M., Chumley, M.J. & Boehm, G.W. (2012) Age exacerbates sickness behavior following exposure to a viral mimetic. Physiol Behav, 105, 1219–1225.

Meinhardt, J., Radke, J., Dittmayer, C., Franz, J., Thomas, C., Mothes, R., Laue, M., Schneider, J., Brunink, S., Greuel, S., Lehmann, M., Hassan, O., Aschman, T., Schumann, E., Chua, R.L., Conrad, C., Eils, R., Stenzel, W., Windgassen, M., Rossler, L., Goebel, H.H., Gelderblom, H.R., Martin, H., Nitsche, A., Schulz-Schaeffer, W.J., Hakroush, S., Winkler, M.S., Tampe, B., Scheibe, F., Kortvelyessy, P., Reinhold, D., Siegmund, B., Kuhl, A.A., Elezkurtaj, S., Horst, D., Oesterhelweg, L., Tsokos, M., Ingold-Heppner, B., Stadelmann, C., Drosten, C., Corman, V.M., Radbruch, H. & Heppner, F.L. (2020) Olfactory transmucosal SARS-CoV-2 invasion as a port of central nervous system entry in individuals with COVID-19. Nat Neurosci.

Meyer, U. (2014) Prenatal poly(i:C) exposure and other developmental immune activation models in rodent systems. Biol Psychiatry, 75, 307–315.

Mian, M.F., Ahmed, A.N., Rad, M., Babaian, A., Bowdish, D. & Ashkar, A.A. (2013) Length of dsRNA (poly I:C) drives distinct innate immune responses, depending on the cell type. J Leukoc Biol, 94, 1025–1036.

Mueller, F.S., Richetto, J., Hayes, L.N., Zambon, A., Pollak, D.D., Sawa, A., Meyer, U. & Weber-Stadlbauer, U. (2019) Influence of poly(I:C) variability on thermoregulation, immune responses and pregnancy outcomes in mouse models of maternal immune activation. Brain Behav Immun, 80, 406–418.

Murray, C., Griffin, E.W., O’Loughlin, E., Lyons, A., Sherwin, E., Ahmed, S., Stevenson, N.J., Harkin, A. & Cunningham, C. (2015) Interdependent and independent roles of type I interferons and IL-6 in innate immune, neuroinflammatory and sickness behaviour responses to systemic poly I:C. Brain, behavior, and immunity, 48, 274–286.

Murray, C., Sanderson, D.J., Barkus, C., Deacon, R.M., Rawlins, J.N., Bannerman, D.M. & Cunningham, C. (2012) Systemic inflammation induces acute working memory deficits in the primed brain: relevance for delirium. Neurobiol Aging, 33, 603–616 e603.

Murray, K.N., Edye, M.E., Manca, M., Vernon, A.C., Oladipo, J.M., Fasolino, V., Harte, M.K., Mason, V., Grayson, B., McHugh, P.C., Knuesel, I., Prinssen, E.P., Hager, R. & Neill, J.C. (2019) Evolution of a maternal immune activation (mIA) model in rats: Early developmental effects. Brain Behav Immun, 75, 48–59.

Palchetti, S., Starace, D., De Cesaris, P., Filippini, A., Ziparo, E. & Riccioli, A. (2015) Transfected poly(I:C) activates different dsRNA receptors, leading to apoptosis or immunoadjuvant response in androgen-independent prostate cancer cells. J Biol Chem, 290, 5470–5483.

Poloni, T.E., Carlos, A.F., Cairati, M., Cutaia, C., Medici, V., Marelli, E., Ferrari, D., Galli, A., Bognetti, P., Davin, A., Cirrincione, A., Ceretti, A., Cereda, C., Ceroni, M., Tronconi, L., Vitali, S. & Guaita, A. (2020) Prevalence and prognostic value of Delirium as the initial presentation of COVID-19 in the elderly with dementia: An Italian retrospective study. EClinicalMedicine, 26, 100490.

Poovassery, J.S., Sarr, D., Smith, G., Nagy, T. & Moore, J.M. (2009) Malaria-induced murine pregnancy failure: distinct roles for IFN-gamma and TNF. J Immunol, 183, 5342–5349.

Richetto, J., Massart, R., Weber-Stadlbauer, U., Szyf, M., Riva, M.A. & Meyer, U. (2017) Genome-wide DNA Methylation Changes in a Mouse Model of Infection-Mediated Neurodevelopmental Disorders. Biol Psychiatry, 81, 265–276.

Shi, L., Fatemi, S.H., Sidwell, R.W. & Patterson, P.H. (2003) Maternal influenza infection causes marked behavioral and pharmacological changes in the offspring. J Neurosci, 23, 297–302.

Shibamiya, A., Hersemeyer, K., Schmidt Woll, T., Sedding, D., Daniel, J.M., Bauer, S., Koyama, T., Preissner, K.T. & Kanse, S.M. (2009) A key role for Toll-like receptor-3 in disrupting the hemostasis balance on endothelial cells. Blood, 113, 714–722.

Skelly, D.T., Griffin, E.W., Murray, C.L., Harney, S., O’Boyle, C., Hennessy, E., Dansereau, M.A., Nazmi, A., Tortorelli, L., Rawlins, J.N., Bannerman, D.M. & Cunningham, C. (2019) Acute transient cognitive dysfunction and acute brain injury induced by systemic inflammation occur by dissociable IL-1-dependent mechanisms. Mol Psychiatry, 24, 1533–1548.

Stetson, D.B. & Medzhitov, R. (2006) Type I interferons in host defense. Immunity, 25, 373–381.

Sun, Q., Sun, L., Liu, H.H., Chen, X., Seth, R.B., Forman, J. & Chen, Z.J. (2006) The specific and essential role of MAVS in antiviral innate immune responses. Immunity, 24, 633–642.

Takeuchi, O., Hemmi, H. & Akira, S. (2004) Interferon response induced by Toll-like receptor signaling. J Endotoxin Res, 10, 252–256.

Taylor, J.M., Minter, M.R., Newman, A.G., Zhang, M., Adlard, P.A. & Crack, P.J. (2014) Type-1 interferon signaling mediates neuro-inflammatory events in models of Alzheimer’s disease. Neurobiology of aging, 35, 1012–1023.

Vabret, N., Britton, G.J., Gruber, C., Hegde, S., Kim, J., Kuksin, M., Levantovsky, R., Malle, L., Moreira, A., Park, M.D., Pia, L., Risson, E., Saffern, M., Salome, B., Esai Selvan, M., Spindler, M.P., Tan, J., van der Heide, V., Gregory, J.K., Alexandropoulos, K., Bhardwaj, N., Brown, B.D., Greenbaum, B., Gumus, Z.H., Homann, D., Horowitz, A., Kamphorst, A.O., Curotto de Lafaille, M.A., Mehandru, S., Merad, M., Samstein, R.M. & Sinai Immunology Review, P. (2020) Immunology of COVID-19: Current State of the Science. Immunity, 52, 910–941.

Varatharaj, A., Thomas, N., Ellul, M.A., Davies, N.W.S., Pollak, T.A., Tenorio, E.L., Sultan, M., Easton, A., Breen, G., Zandi, M., Coles, J.P., Manji, H., Al-Shahi Salman, R., Menon, D.K., Nicholson, T.R., Benjamin, L.A., Carson, A., Smith, C., Turner, M.R., Solomon, T., Kneen, R., Pett, S.L., Galea, I., Thomas, R.H., Michael, B.D. & CoroNerve Study, G. (2020) Neurological and neuropsychiatric complications of COVID-19 in 153 patients: a UK-wide surveillance study. Lancet Psychiatry.

Wang, J., Campbell, I.L. & Zhang, H. (2008) Systemic interferon-alpha regulates interferon-stimulated genes in the central nervous system. Mol Psychiatry, 13, 293–301.

Weber, F., Wagner, V., Rasmussen, S.B., Hartmann, R. & Paludan, S.R. (2006) Double-stranded RNA is produced by positive-strand RNA viruses and DNA viruses but not in detectable amounts by negative-strand RNA viruses. J Virol, 80, 5059–5064.

Weintraub, M.K., Kranjac, D., Eimerbrink, M.J., Pearson, S.J., Vinson, B.T., Patel, J., Summers, W.M., Parnell, T.B., Boehm, G.W. & Chumley, M.J. (2014) Peripheral administration of poly I:C leads to increased hippocampal amyloid-beta and cognitive deficits in a non-transgenic mouse. Behav Brain Res, 266, 183–187.

Zhou, Y., Guo, M., Wang, X., Li, J., Wang, Y., Ye, L., Dai, M., Zhou, L., Persidsky, Y. & Ho, W. (2013) TLR3 activation efficiency by high or low molecular mass poly I:C. Innate Immun, 19, 184–192.

Zhu, X., Levasseur, P.R., Michaelis, K.A., Burfeind, K.G. & Marks, D.L. (2016) A distinct brain pathway links viral RNA exposure to sickness behavior. Sci Rep, 6, 29885.

